# Relating local connectivity and global dynamics in recurrent excitatory-inhibitory networks

**DOI:** 10.1101/2022.08.25.505122

**Authors:** Yuxiu Shao, Srdjan Ostojic

## Abstract

How the connectivity of cortical networks determines the neural dynamics and the resulting computations is one of the key questions in neuroscience. Previous works have pursued two complementary strategies to quantify the structure in connectivity, by specifying either the local statistics of connectivity motifs between small groups of neurons, or by defining network-wide low-rank patterns of connectivity that determine the resulting low-dimensional dynamics. A direct relationship between these two approaches is however currently missing, and in particular it remains to be clarified how local connectivity statistics are related to the global connectivity structure and shape the low-dimensional activity. To bridge this gap, here we develop a method for mapping local connectivity statistics onto an approximate global low-rank structure. Our method rests on approximating the global connectivity matrix using dominant eigenvectors, which we compute using perturbation theory for random matrices. This approach demonstrates that multi-population networks defined from local connectivity properties can in general be approximated by low-rank connectivity with Gaussian-mixture statistics. We specifically apply this method to excitatory-inhibitory networks, and show that it leads to accurate predictions for both the low-dimensional dynamics, and for the activity of individual neurons. Altogether, our approach allows us to disentangle the effects of mean connectivity and reciprocal motifs on the global recurrent feedback, and provides an intuitive picture of how local connectivity shapes global network dynamics.

**Author summary:** The structure of connections between neurons is believed to determine how cortical networks control behaviour. Current experimental methods typically measure connections between small numbers of simultaneously recorded neurons, and thereby provide information on statistics of local connectivity motifs. Collective network dynamics are however determined by network-wide patterns of connections. How these global patterns are related to local connectivity statistics and shape the dynamics is an open question that we address in this study. Starting from networks defined in terms of local statistics, we develop a method for approximating the resulting connectivity by global low-rank patterns. We apply this method to classical excitatory-inhibitory networks and show that it allows us to predict both collective and single-neuron activity. More generally, our approach provides a link between local connectivity statistics and global network dynamics.

## Introduction

One of the central questions in neuroscience is how the connectivity structure of cortical networks determines the collective dynamics of neural activity and their function. Experimental assessments of connectivity are typically based on measurements of synaptic weights between small numbers of neurons recorded simultaneously [1–9]. The most common approach to quantify connectivity therefore focuses on *local statistics*, and starts by characterizing the connection probability between pairs of neurons based on their type, before considering progressively more complex connectivity motifs. Linking these local connectivity statistics to the emerging network dynamics has been an active topic of investigations [10–24]. A second approach, motivated by computational network models instead of experimental measurements [25–29], instead specifies the connectivity in terms of a low-rank structure defined by network-wide patterns of connectivity [30–39]. This global connectivity structure directly determines the low-dimensional dynamics and resulting computations [30, 31, 33], yet it remains unclear how it is related to local connectivity statistics that can be recorded experimentally. In this study, we aim to bridge this gap, by mapping local connectivity statistics onto a global, low-rank description of connectivity and comparing the resulting dynamics.

Starting from random networks with connectivity defined in terms of local, cell-type dependent statistics, we develop a low-rank approximation based on the dominant eigenmodes of the connectivity matrix. Using perturbation theory, we show that the obtained low-rank connectivity patterns universally obey Gaussian-mixture statistics and therefore lead to analytically tractable dynamics [31, 33]. We specifically apply this approach to excitatory-inhibitory networks with connections consisting of independent and reciprocal parts, and exploit the low-rank approximation to predict the emerging dynamics.

We first show that, although the dominant low-rank structure is set on average by the mean synaptic weights [40–42], a perturbative approach accurately predicts the components of individual neurons on the dominant eigenvectors for individual instances of the random connectivity. As a result, our low-rank approximation analytically predicts the activity of individual neurons in the original E-I network defined based on local statistics. The analytic description of the dynamics in the low-rank approximation moreover leads to the identification of two distinct sources of recurrent feedback corresponding respectively to the mean connectivity and reciprocal connections between neurons. In particular, the reciprocal motifs impact dynamics by modulating both the dominant eigenvalue and the corresponding eigenvectors, and can give rise to additional bistability in the network. Altogether, our analytical mapping of the local EI statistics to a low-rank description provides a quantitative and intuitive description of how local connectivity statistics determine global low-dimensional dynamics.

## 1 Results

### 1.1 Local vs. global representations of random recurrent connectivity

We study networks of *N* rate units with random recurrent connectivity given by the connectivity matrix **J**, where the entry *J*_*ij*_ corresponds to the strength of the synapse from neuron *j* to neuron *i*. A full statistical description of the random connectivity would require specifying the joint distribution *P* ({*J*_*ij*_}) of the *N* ^2^ synaptic weights. Determining the dynamics from this high-dimensional distribution is however in general intractable. We therefore focus on connectivity models that make simplifying assumptions on the underlying statistics.

Our specific goal is to relate two different classes of such models, which we refer to as the *local* and the *global* representations of recurrent connectivity. Both representations assume that the network consists of *P* populations, and the statistics of connectivity depend only on the pre- and post-synaptic populations. The two representations however take as starting points different statistical features of the connectivity.

The local representation defines the connectivity statistics by starting from the marginal distributions Prob(*J*_*ij*_ = *J*) of individual synaptic weights, and by including progressively higher-order correlations referred to as *connectivity motifs* [1, 10, 14]. In this work, we will consider only the first two orders, i. e. the distribution of individual weights and the pairwise correlations *η*_*ij*_ between reciprocal connections *J*_*ij*_ and *J*_*ji*_ that quantify pairwise motifs (Fig 1A). Our key assumption is that both the marginal distributions of *J*_*ij*_ and the correlations *η*_*ij*_ depend only on the populations *p* and *q* that the post- and pre-synaptic neurons belong to:

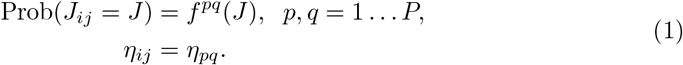

**Fig 1.**
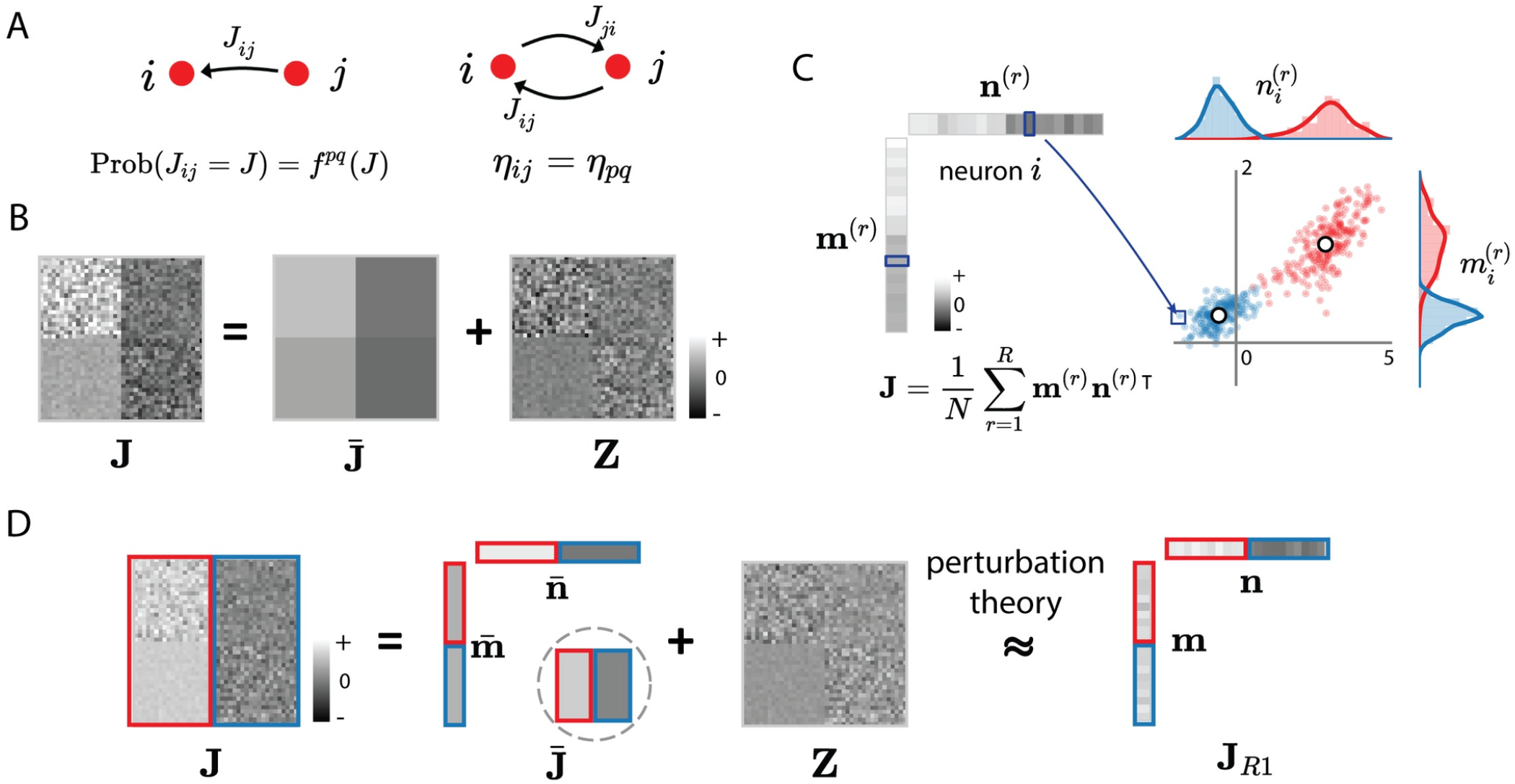
Local vs global representations of recurrent connectivity. (A) The local representation defines the statistics of synaptic weights *J*_*ij*_ by starting from the marginal probability distribution of individual synaptic weights (left) and then specifying reciprocal motifs in terms of correlations *η*_*ij*_ between reciprocal weights *J*_*ij*_ and *J*_*ji*_ connecting neurons *i* and *j* (right). Both the marginal distribution and the reciprocal correlations are assumed to depend only on the populations *p* and *q* that the neurons *i* and *j* belong to. (B) The resulting connectivity matrix **J** has block-structured statistics, where different blocks correspond to connections between the *P* different populations (*P* = 2 in this illustration). It can be decomposed into a superposition of a mean component 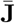, and a remaining zero-mean random connectivity component **Z** that has block-structured variances. (C) The global, low-rank representation defines the connectivity matrix **J** as the sum of *R* outer products between connectivity vectors **m**^(*r*)^, **n**^(*r*)^ for *r* = 1 … *R*. The statistics of connectivity are defined in terms of the joint probability distribution over neurons *i* of their entries 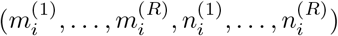 on connectivity vectors. We specifically consider the class of Gaussian-mixture low-rank models, where each neuron is first assigned to a population *p*, and within each population the entries on connectivity vectors are generated from a multivariate Gaussian distribution with fixed statistics. Here we illustrate this distribution for one pair of connectivity vectors (*R* = 1) and *P* = 2 populations. Each dot represents the connectivity parameters 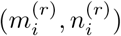 of one neuron *i*, the red and blue colours denote the two populations, white dots and the rotations of the dot clouds indicate the mean and covariance of the distribution for each population. (D) Relating the local and global representations of recurrent connectivity for a simplified excitatory-inhibitory network. In this model, the mean connectivity depends only on the presynaptic population (indicated by red and blue colours). The mean connectivity 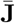 is in this case rank-one, and can be written as an outer product of vectors 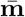 and 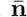. We approximate the full connectivity by a rank-one matrix, with connectivity vectors **m** and **n** obtained from 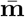 and 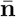 using perturbation theory.

All synapses connecting the same two populations therefore have identical statistics, leading to a block-like statistical structure for the connectivity matrix **J** (Fig 1B left panel).

The global representation of connectivity instead refers to the situation where **J** is defined as a low-rank matrix [30, 31, 33]:

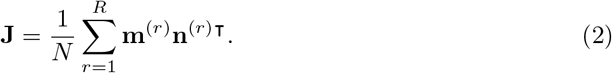

Here 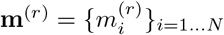 and 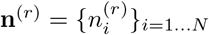 for *r* = 1 … *R* are referred to as *connectivity vectors*, where *R* is the rank of **J**. In this representation, the statistics of connectivity are defined by the distribution of vector elements, rather than directly by the distribution of synaptic weights as in the local representation. Specifically, each neuron *i* is characterized by its set of entries 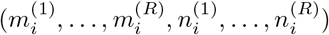 over the connectivity vectors. For each neuron, these 2*R* entries are generated from a joint distribution, independently of the other neurons, and the parameters of this joint distribution depend on the population *p* the neuron belongs to. Here we focus on the broad class of Gaussian-mixture low-rank networks, in which for population *p*, the joint distribution of elements is a multi-variate Gaussian defined by the means and covariances of the 2*R* entries [31, 33] (Fig 1C).

To relate the local and the global representations of connectivity, a key observation is that any matrix **J** generated from the local statistics defined in Eqs. (1) can be expressed as

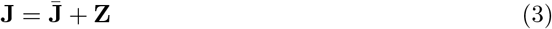

where 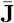 contains the mean values of the connections, and **Z** contains the remaining, zero-mean random part [40]. Because of the underlying population structure (Eqs. (1)), 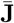 consists of *P*×*P* blocks with identical values within each block (Fig 1B middle panel), and is therefore at most of rank *P*. The random part **Z** is instead in general of rank *N*, but obeys block-like statistics, with variance and normalized covariance parameters defined by *P*×*P* matrices (Methods Secs. 2.1.1-2.1.2, Eqs. (25), (30)).

For the sake of simplicity, in this study, we focus on a simplified excitatory-inhibitory model [41, 43]. This network consists of one excitatory and one inhibitory population, so *P* = 2 and in the following we use the population indices *p, q* = *E, I*. A central simplifying assumption in this model is that the mean synaptic weights depend only on the pre-synaptic population, so that 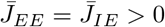 and 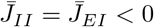. The mean connectivity matrix 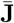 therefore consists of only two blocks and is unit rank (Fig 1D). The statistics of the random part **Z** instead depend on both pre- and post-synaptic populations, and are therefore described by 2×2 matrices of variance and normalized covariance parameters (see Methods Sec. 2.1.3, Eq. (35)).

### 1.2 Approximating locally-defined connectivity with low-rank connectivity

To relate the local and global representations of connectivity, we start from a connectivity matrix **J** generated from the local statistics (Eqs. (1)) and approximate it by a rank-*R* matrix of the form given in Eq. (2). As the locally-defined connectivity matrix **J** is of rank *N*, this is equivalent to the classical low-rank approximation problem, for which a variety of methods exist [30, 31, 33]. Here we use simple truncated eigen-decomposition as it preserves the dominant eigenvalues that determine non-linear dynamics.

Applying the standard eigenmode decomposition, **J** can be in general factored as

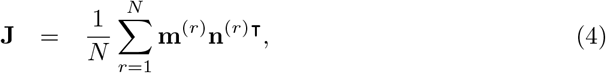

where **m**^(*r*)^ and **n**^(*r*)^ are rescaled versions of the *r*-th right and left eigenvectors (Methods Sec. 2.3, Eqs (42)-(47)), ordered by the absolute value of their eigenvalue *λ*_*r*_ for *r* = 1 … *N*. A rank-*R* approximation that preserves the top *R* eigenvalues can then be obtained by simply keeping the first *R* terms in the sum in Eq. (4). In this study, we focus on *R* = 1, corresponding to the dominant eigenvalue. Higher order approximations will be described elsewhere.

Eigenvalues and eigenvectors are in general complex non-linear functions of the entries of the matrix **J**. To determine the dominant eigenvalues and the corresponding vectors of **J**, we capitalize on the observation in Eq. (3) that a locally-defined connectivity matrix can in general be expressed as a sum of a low-rank matrix of mean values 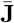 and the remaining random part **Z**. Previous studies have found that the eigenspectra of matrices with such structure typically consist of two components in the complex plane: a continuously-distributed bulk determined by the random part, and discrete outliers controlled by the low-rank structure [30, 32, 39, 44–46]. In this study, we extend previous approaches to determine the influence of the block-like statistics of **Z** on the outliers that correspond to dominant eigenvalues. We then use perturbation theory to determine the corresponding left and right eigenvectors and their statistical structure. Here we summarize the main steps of this analysis (full details are provided in Methods), and then apply it to specific cases in the following sections.

We focus on the simplified E-I network for which the mean part of the connectivity is unit rank and can therefore be written as 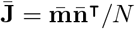, so that the full connectivity matrix is

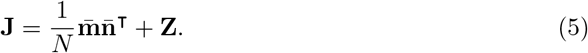

The mean part 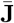 of the connectivity has a unique non-trivial eigenvalue 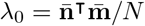 which can give rise to one or several outliers *λ* in the eigenspectrum of **J**. To determine how the random part of the connectivity influences *λ*, we start from the characteristic equation for the eigenvalues of **J** and exploit Eq. (5) to apply the matrix determinant lemma (Eq. (57)). This leads to a non-linear equation for *λ* [32]:

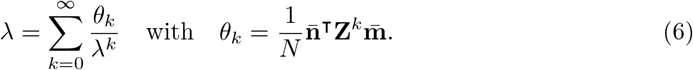

Truncating the sum to second order yields an approximate third order polynomial for *λ*:

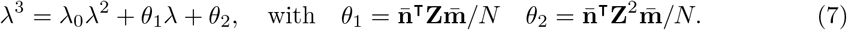

The statistics of the outlying eigenvalue can then be obtained by averaging over the random part of the connectivity **Z**.

An approximate expression for the right and left connectivity vectors **m** and **n** of **J** corresponding to the outliers *λ* can be determined using first order perturbation theory [47]. We first note that 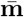 and 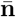 are the right- and left-eigenvectors of 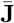 corresponding to the non-trivial eigenvalue *λ*_0_. Interpreting the full connectivity matrix **J** as 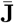 perturbed by a random matrix **Z**, at first order **m** and **n** can be expressed as

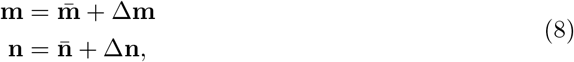

with

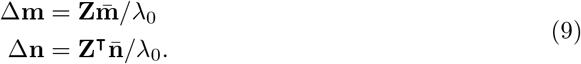

A key observation is that each element of Δ**m** and Δ**n** is a sum of *N* random variables. The central limit theorem therefore predicts that, in the limit of large *N*, the statistics of Δ*m*_*i*_ and Δ*n*_*i*_, and therefore *m*_*i*_ and *n*_*i*_, follow a Gaussian distribution. In general, the mean and variance of *m*_*i*_ and *n*_*i*_ and their correlation are determined by the mean, variance and correlation of the elements of **Z**, but not the specific form of the probability distribution. Since the matrix **Z** has block-like statistics determined by the population structure, the statistics of the resulting *m*_*i*_ and *n*_*i*_ depend on the population *p* the neuron *i* belongs to. Overall, the distribution of elements of **m** and **n** obtained from perturbation theory therefore follow a Gaussian-mixture distribution, so that our approach effectively approximates a locally-defined **J** by a Gaussian-mixture low-rank model specified by the means 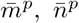 the variances 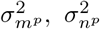 and the covariances 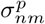 of the entries on the connectivity vectors for *p* = *E, I*.

We next apply the perturbative approach described here to networks with independent random components, and then to networks with reciprocal motifs.

### 1.3 Low-rank structure induced by independently generated synaptic connections

We first apply our approach for a low-rank approximation to the simplest version of the locally-defined excitatory-inhibitory network where each *J*_*ij*_ is generated independently from a Gaussian distribution with a mean that depends only on the pre-synaptic population, i. e. 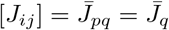 with *p, q* ∈ *E, I*. The entries of the eigenvectors 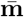 and 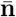 of the mean connectivity matrix 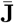 are then given by:

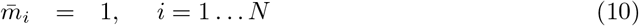

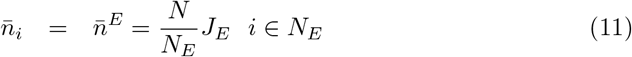

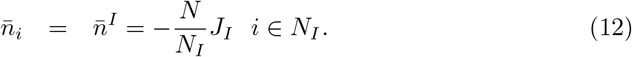

We first consider the case where the variance of *J*_*ij*_ is uniform across all connections and given by *g*^2^*/N*. In that situation, the entries of the random part of the connectivity **Z** are independent, identically distributed Gaussians, and the eigenvalue spectrum of **Z** converges to a uniform distribution on the complex plane within a circle of radius *g* [48–50]. Previous studies [30, 32, 44, 45] have shown that adding a unit rank matrix on top of an i. i. d. random matrix as in Eq. (3) leads to an eigenspectrum that is the superposition of the two spectra, and therefore consists of a circular bulk of radius *g* and an outlier on average located at the eigenvalue *λ*_0_ of 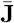. Our analyses of the eigenvalues of **J** confirm this result (Fig 2A). Indeed, averaging over **Z** in Eq. (6), [*θ*_*k*_] = 0 for all *k* [32], so that the outlier is on average given by [*λ*] = *λ*_0_. Our approach moreover gives an expression for the standard deviation of the outlier which grows linearly with *g* (Fig 2B, Eq. (78)). Examining the entries of the left and right eigenvectors **n** and **m** of **J** corresponding to the outlier, we find that perturbation theory accurately predicts the individual entries of the eigenvectors as long as *g* is sufficiently below unity (Fig 2D). As expected, the distribution of (*m*_*i*_, *n*_*i*_) is well described by a mixture of two Gaussians centred at 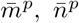. Perturbation theory provides a lower bound for the values of the corresponding variances (Fig 2E). For large values of *g*, the distributions remain Gaussian, but their variances increase above the predictions of perturbation theory. Importantly, the entries of the left and right eigenvectors are uncorrelated, and only their means, but not their variances, differ between the two populations (Figs 2C, E).

**Fig 2.**
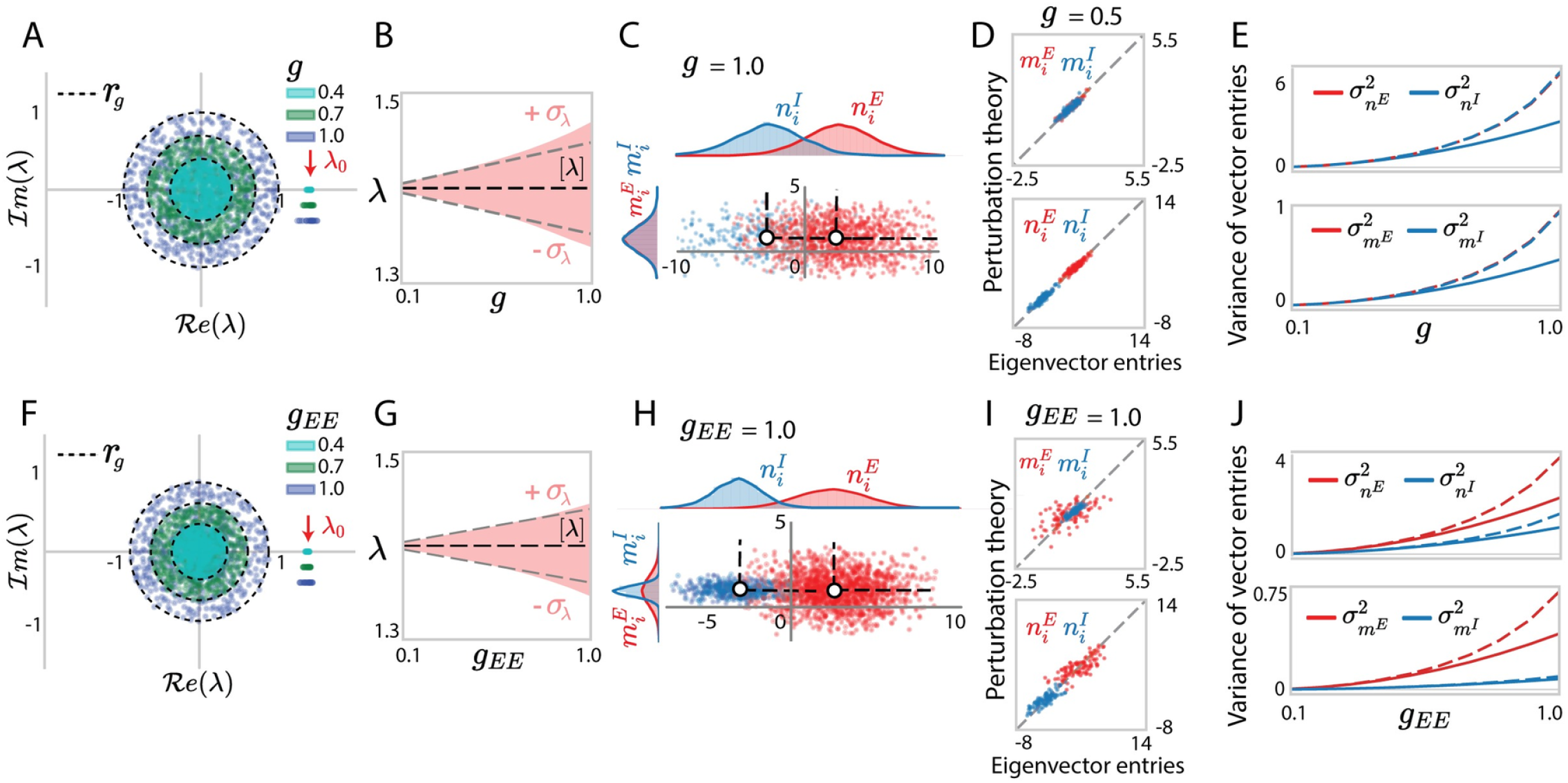
Eigenvalues and dominant eigenvectors for locally-defined Gaussian connectivity with independent synaptic weights. (A) Eigenvalue spectra of excitatory-inhibitory connectivity matrices **J** with elements generated from Gaussian distributions with identical variances *g*^2^*/N* over neurons. The coloured dots in the circular bulk shows 600 eigenvalues for one realization of the random connectivity for each value of *g*. Different colours correspond to different values of *g*. Dashed envelopes indicate the theoretical predictions for the radius *r*_*g*_ of the circular bulk computed according to Eqs. (146), (147). Outlying eigenvalues are shown for 30 realizations of the random connectivity, and for different *g* their location on the *y*-axis is shifted to help visualization. The red arrow points to the eigenvalue *λ*_0_ of the mean connectivity matrix 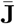. (B) Statistics of outlying eigenvalues over realizations of random connectivity. Empirical distribution (the red area shows mean ± standard deviation), compared with the theoretical predictions for the mean (black dashed line) and standard deviation (gray dashed line) obtained using Eq. (78). (C) Scatter plot showing for each neuron *i* its entry *n*_*i*_ on the left eigenvector against its entry *m*_*i*_ on the right eigenvector. Red and blue colours represent respectively excitatory and inhibitory neurons. The white dots and the dashed lines respectively indicate the means and covariances for each population obtained from simulations. (D) Comparison between eigenvector entries obtained from direct eigen-decomposition of **J** with predictions of perturbation theory (Eqs. (8), (9)). (E) Comparison between simulations (full lines) and theory (dashed lines) for the variances 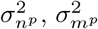 of eigenvector entries corresponding to different populations (Eq. (88)). (F-J) Identical quantities for connectivity matrices in which the variance parameters are heterogeneous: *g*_*EE*_ : *g*_*EI*_ : *g*_*IE*_ : *g*_*II*_ = 1.0 : 0.5 : 0.2 : 0.8, *g*_*EE*_ increases from 0 to 1. Other network parameters *N*_*E*_ = 4*N*_*I*_ = 1200 and *J*_*E*_ = 2.0, *J*_*I*_ = 0.6 in all simulations.

We next turned to the case where the variances of synaptic weights depend on the pre- and post-synaptic populations *q, p*, and are given by 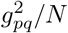. In that case, the entries of the random part of the connectivity **Z** are independent, but not identically distributed Gaussians. Previous studies [11, 44] have shown that the spectrum of **Z** remains circularly symmetric, but its radius *r*_*g*_ is determined by a combination of variance parameters *g*_*pq*_ (S4 Appendix, Eqs. (146), (147)). Examining the resulting connectivity matrix **J**, we found that the results for the uniform case directly extend to this heterogeneous situation. The eigenspectrum of **J** still consists of an independent superposition of the spectra of **Z** and 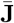 (Fig 2F). In particular, the random part of the connectivity does not modify the average value of the outlier, but only impacts its variance, which now depends on a combination of the variances *g*_*pq*_ (Fig 2G, Eq. (88)). Similarly to the uniform case, the distribution of the entries of the left and right eigenvectors is well described by a mixture of two Gaussians, with variances predicted by perturbation theory (Fig 2I). The entries of the left and right eigenvectors are uncorrelated, but now both their means and variances depend on the population the neuron belongs to (Figs 2H, J).

In summary, when synaptic connections *J*_*ij*_ are generated independently across pairs of neurons, the equivalent global representation is a Gaussian-mixture low-rank model where the entries of the structure vectors are independent with mean values determined by the low-rank structure of the mean connectivity matrix 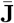. Importantly, in that situation, the dominant outlying eigenvalues of **J** are on average identical to those of 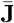.

### 1.4 Low-rank structure induced by reciprocal motifs

We next turn to locally-defined excitatory-inhibitory networks with reciprocal connectivity motifs quantified by the correlation *η*_*ij*_ between reciprocal synaptic weights *J*_*ij*_ and *J*_*ji*_. We assumed that these reciprocal correlations are identical for any pair of neurons *i* and *j* belonging to a given pair of populations *p* and *q*, and used the corresponding parameters *η*_*pq*_ to generate the connectivity matrix **J** (Methods Sec. 2.1.2). Within the decomposition of **J** in a mean 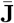 and random part **Z** (Eq. (22)), the additional reciprocal correlations affect only the statistics of **Z**.

We first consider the homogeneous case where the reciprocal correlation is identical across all populations, i. e. *η*_*pq*_ = *η*. Previous studies have shown that a random matrix **Z** with zero mean and reciprocal correlations *η* has a continuous spectrum that is deformed from a circle into an ellipse as *η* is increased [18, 51]. Superpositions between correlated random matrices, and low-rank structure such as 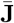 have to our knowledge not been previously studied. Inspecting the eigenspectrum of 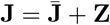, we found that it still consists of a continuous bulk and discrete outliers (Fig 3A). The continuous bulk is contained in an ellipse in the complex plane identical to the spectrum of **Z**, as in the uncorrelated case. In contrast, we found that the outliers deviated from the eigenvalues of 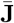 as *η* was increased. These deviations were well captured by our analytic approach summarized in Eq. (7). Indeed, when averaging Eq. (7) over **Z**, reciprocal correlations generate a non-zero 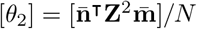 due to **Z**^2^. This term leads to a cubic equation in Eq. (7) and therefore has two potential effects. First, the non-zero *θ*_2_ induces deviations of the outliers from the eigenvalue *λ*_0_ of 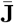. The direction of these deviations is positive if excitation dominates (*λ*_0_ = *J*_*E*_ − *J*_*I*_ *>* 0) and negative if inhibition dominates (*λ*_0_ = *J*_*E*_ −*J*_*I*_ *<* 0, Figs 4F-H). Second, the cubic equation can have up to three solutions and therefore potentially generates additional outliers, and in particular complex conjugate ones (Fig 3M). Whether these additional outliers are observed depends on the accuracy of the third-order approximation to the determinant lemma, and on the norm of these outliers compared to the spectral radius (Fig 4E).

**Fig 3.**
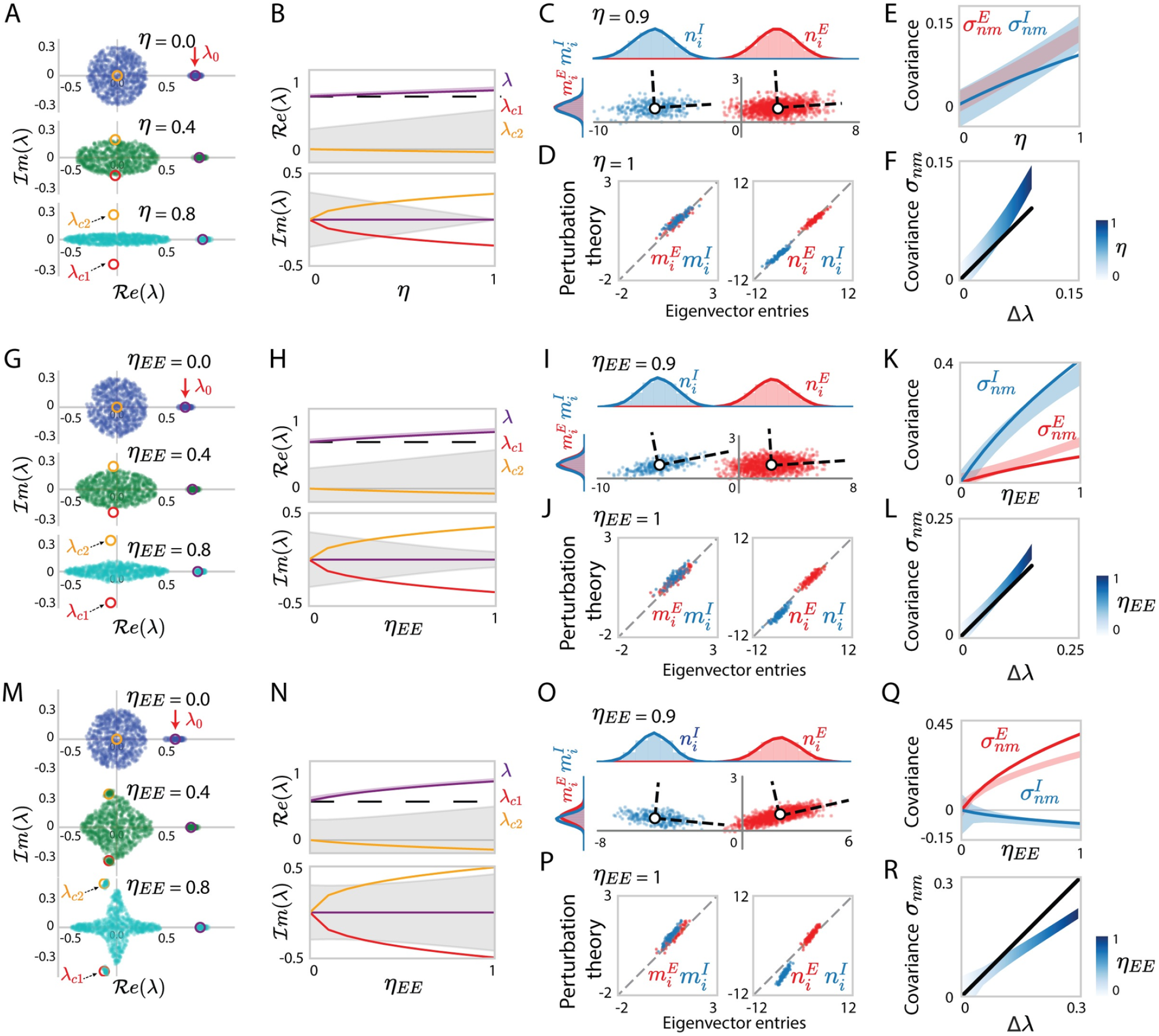
Eigenvalues and dominant eigenvectors for locally-defined Gaussian connectivity with reciprocal motifs. (A) Eigenvalue spectra of excitatory-inhibitory connectivity matrices **J**, with homogeneous reciprocal correlations *η*. Different colours correspond to networks with different values of *η*. The dots in the elliptical bulk show 600 eigenvalues for one realization of the random connectivity. Outlying eigenvalues are shown for 30 realizations of the random connectivity. The red arrow on the top points to the eigenvalue *λ*_0_ of the mean connectivity 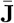. Coloured circles are the eigenvalues predicted using determinant lemma (Eq. (7)). (B) Comparison of the eigenvalues from the finite-size simulation with the predictions of the determinant lemma as the reciprocal correlation *η* is increased. The coloured solid lines show the roots of the third-order polynomial in Eq. (7). The light purple area indicates the empirical distribution of the dominant outlier, while the black dashed line is the unperturbed eigenvalue *λ*_0_. The grey areas represent the areas covered by the eigenvalue bulk. (C) Scatter plot showing for each neuron *i* its entry *n*_*i*_ on the left eigenvector against its entry *m*_*i*_ on the right eigenvector. Red and blue colours represent respectively excitatory and inhibitory neurons. The white dots and the dashed lines respectively indicate the means and covariances for each population. (D) Comparison between eigenvector entries obtained from direct eigen-decomposition of **J** with predictions of perturbation theory (Eqs. (8), (9)). (E) Comparison between simulations (coloured areas) and theoretical predictions (coloured lines, Eq. (95)) for the population covariance 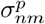 of the entries on the left and right connectivity eigenvectors to different populations. (F) Comparison of the overall covariance *σ*_*nm*_ (Eq. (70)) with the deviation Δ*λ* of the dominant outlying eigenvalue from the unperturbed value *λ*_0_. Empirical covariance (gradient blue area, colour depth represents *η*) compared with the theoretical prediction (black line) obtained using Eqs. (95), (90). The *x*-axis uses the theoretical prediction of the deviation of the eigenvalue *λ* from *λ*_0_. (G-L) Same as (A-F) for a connectivity matrix with heterogeneous reciprocal correlations: *η*_*EE*_ = *η*_*EI*_ =−*η*_*II*_ *>* 0, and *η*_*EE*_ increasing from 0 to 1. (M-R) Same as (A-F) for a connectivity matrix with heterogeneous reciprocal correlations: *η*_*EE*_ = −*η*_*EI*_ = −*η*_*II*_ *>* 0, and *η*_*EE*_ increasing from 0 to 1. Other network parameters: *N*_*E*_ = 4*N*_*I*_ = 1200 and homogeneous variance parameters *g*_*pq*_ = *g* = 0.3 in all simulations, *J*_*E*_ = 2.0, *J*_*I*_ = 1.2 for networks in (A-F), *J*_*E*_ = 2.0, *J*_*I*_ = 1.3 for networks in (G-L), *J*_*E*_ = 2.0, *J*_*I*_ = 1.4 for networks in (M-R).

**Fig 4.**
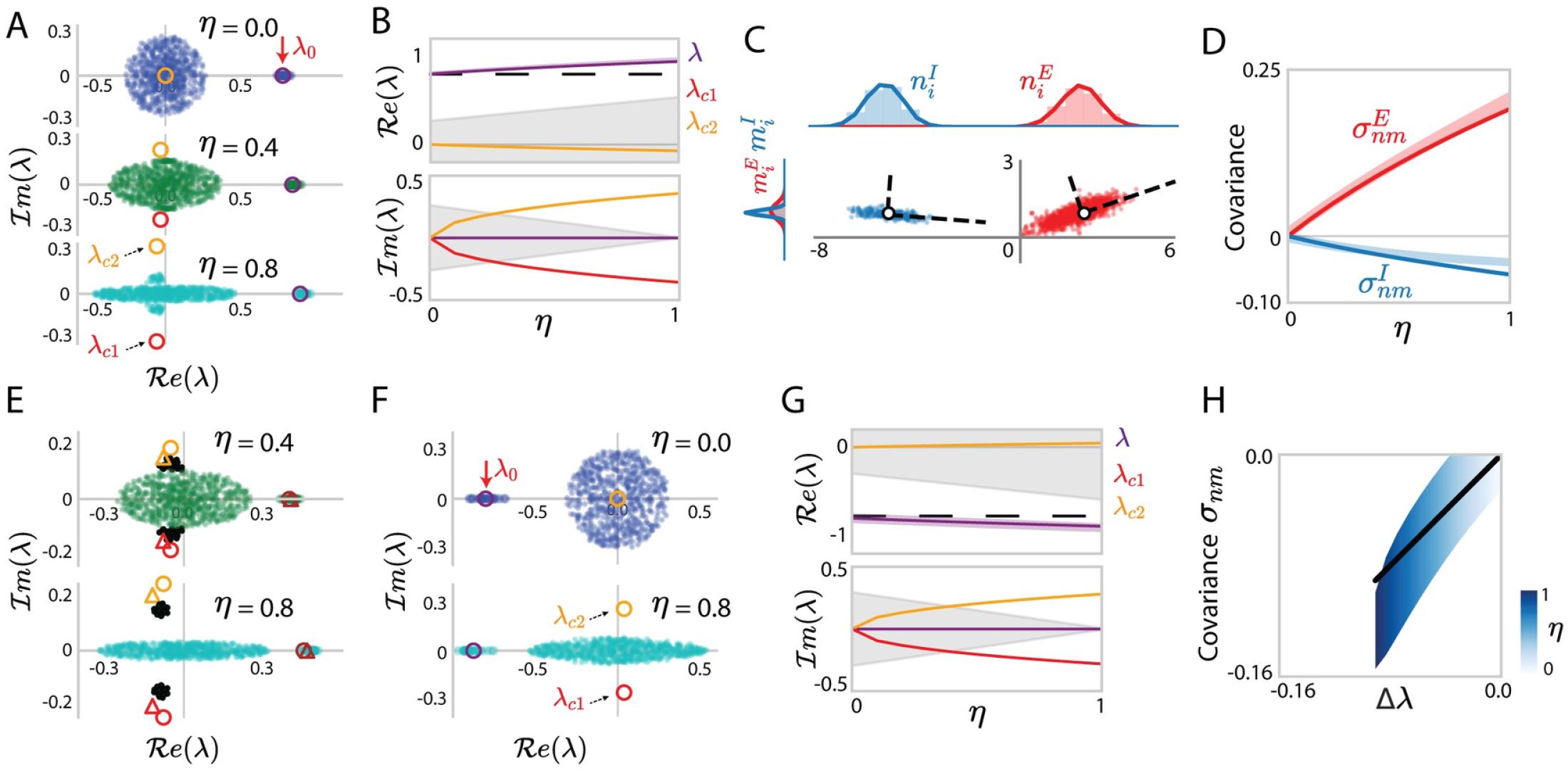
Eigenvalues and dominant eigenvectors for different network connectivity with reciprocal motifs. (A) Eigenvalue spectra of excitatory-inhibitory connectivity matrices **J**, with homogeneous reciprocal correlations *η* but cell-type-dependent variance parameters *g*_*EE*_ : *g*_*EI*_ : *g*_*IE*_ : *g*_*II*_ = 1.0 : 0.5 : 0.2 : 0.8 and *g*_*EE*_ = 0.3. Different colours correspond to networks with different *η*. The elliptical bulk shows 600 randomly sampled eigenvalues for one realization of the random connectivity. Outlying eigenvalues are shown for 30 realizations of the random connectivity. The red arrow on the top points to the unperturbed eigenvalue *λ*_0_ of the mean connectivity 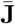. Coloured circles are the eigenvalues predicted using determinant lemma. (B) Comparison of the eigenvalues from the finite-size simulation with the predictions of the determinant lemma as the reciprocal correlation *η* is increased. The coloured solid lines show the roots of the third-order polynomial in Eq. (7). The light purple area indicates the empirical distribution of the dominant outlier, while the black dashed line is the unperturbed eigenvalue *λ*_0_. The grey areas represent the eigenvalue bulk. (C) Scatter plot showing for each neuron *i* its entry *n*_*i*_ on the left eigenvector against its entry *m*_*i*_ on the right eigenvector, with the eigenvectors corresponding to eigenvalue outliers *λ* that deviate from *λ*_0_. Red and blue colours represent respectively excitatory and inhibitory neurons. The white dots and the dashed lines respectively indicate the means and covariances for each population. (D) Comparison between the population covariance 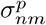 of the entries on the left and right connectivity eigenvectors to different populations (coloured areas) and the predictions of perturbation theory (coloured lines, Eq. (95)). Other network parameters for (A-D): *J*_*E*_ = 2.0, *J*_*I*_ = 1.2, *N*_*E*_ = 4*N*_*I*_ = 1200. (E) Same as (A) for a connectivity matrix with homogeneous reciprocal correlations *η*, the coloured circles are solutions considering *k* up to 2 and the coloured triangles are solutions considering *k* up to 4. Other network parameters: *J*_*E*_ = 1.5, *J*_*I*_ = 1.2, *g*_*EE*_ : *g*_*EI*_ : *g*_*IE*_ : *g*_*II*_ = 1.0, 0.5, 0.2, 0.8, *g*_*EE*_ = 0.2 and *N*_*E*_ = 4*N*_*I*_ = 1200. (F, G) Same as (A, B) for an inhibition dominates connectivity matrix where *J*_*I*_ = 2.0, *J*_*E*_ = 1.2, with homogeneous reciprocity *η* and variance parameters *g* = 0.3. (H) Comparison of the overall covariance *σ*_*nm*_ with the deviation Δ*λ* of the dominant outlying eigenvalue from the unperturbed value *λ*_0_.

We next examine the right- and left-eigenvectors **m** and **n** corresponding to the dominant outlier. Analogous to the uncorrelated case in the networks with independent connections, the individual entries of these vectors are accurately predicted by perturbation theory and exhibit Gaussian-mixture statistics (Fig 3D). Unlike in the uncorrelated case, reciprocal correlations now induce correlations between Δ*m*_*i*_ and Δ*n*_*i*_ (Fig 3C). Indeed, perturbation theory predicts that the first-order effects Δ**m** and Δ**n** of the random connectivity on **m** and **n** are respectively determined by **Z** and its transpose **Z**^⊤^ (Eq. (9)). Reciprocal correlations between *z*_*ij*_ and *z*_*ji*_ directly lead to correlations between **Z** and **Z**^⊤^ and therefore a non-zero covariance *σ*_*nm*_ between elements of **m** and **n**, that can be predicted by mean field theory (Fig 3E, Eq. (70)). When the network has both homogeneous variance parameters and correlation parameters, the excitatory and inhibitory populations have the same covariance 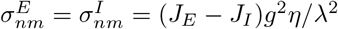 (Figs 3C, E, Eq. (95)). If the synaptic variances *g*_*pq*_ differ across populations, the covariances *σ*_*nm*_ are different for excitatory and inhibitory populations even if the reciprocal correlations are uniform (Figs 4C, D). The strength of the overall covariance reflects the strength of the additional feedback loop due to reciprocal correlations, and is therefore directly related to the deviations of the outlying eigenvalue from the uncorrelated value *λ*_0_ (Fig 3F).

These results directly extend to networks with heterogeneous reciprocal correlations *η*_*pq*_, *p, q* = *E, I*. In particular, in this case finite-size simulations confirm the presence of additional, complex conjugate outliers predicted by the cubic term in Eq. (7) (coloured circles containing outlier scatters at conjugate positions in Fig 3M). Moreover, the covariances 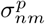 between the entries of low-rank connectivity vectors in this case differ between the excitatory and inhibitory population (Figs 3I, K, O, Q).

### 1.5 Approximating low-dimensional dynamics for locally-defined connectivity

In previous sections, we developed a rank-one approximation of locally-defined excitatory-inhibitory connectivity. Here we use this approximation to describe the resulting low-dimensional dynamics. We consider networks of rate units, where the activation *x*_*i*_ of unit *i* obeys

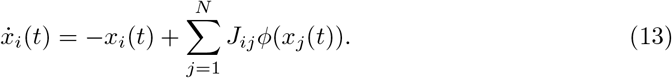

Here *ϕ*(*x*) = 1 + tanh (*x − θ*) is a positive transfer function, and for simplicity, we focus on autonomous dynamics without external inputs. We start from a locally-defined excitatory-inhibitory connectivity matrix, and compare the resulting activity with the theoretical predictions of our rank-one approximation, for which the dynamics are low-dimensional and analytically tractable. We first summarize the theoretical predictions for those dynamics, and then examine the specific cases of independent and reciprocally-correlated connectivity.

Recent works have showed that in networks with a rank *R* connectivity matrix, the trajectories **x**(*t*) ={*x*_*i*_(*t*) }_*i*=1…*N*_ are confined to a low-dimensional subspace of the *N*–dimensional space describing the activity of all units [30–33]. In absence of external inputs, this subspace is *R*-dimensional and spanned by the set of connectivity eigenvectors **m**^(*r*)^ for *r* = 1 … *R*, so that the trajectories can be parametrized as 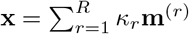 where *κ*_*r*_ is a collective latent variable representing activity along **m**^(*r*)^. For a rank-one (*R* = 1) connectivity corresponding to an approximation of our locally-defined E-I network, the dynamics can therefore be represented by a single latent variable *κ*, so that the activation of unit *i* is given by

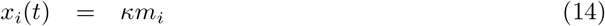

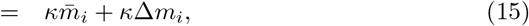

where we inserted the expression for *m*_*i*_ obtained from a first-order perturbation (Eq. (8), (9)). Note that since 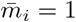, the first term in the r. h. s. of Eq. (15) corresponds to population-averaged activity, while the second term is the deviation of the activity of unit *i* from the population average. Inserting the values for Δ*m*_*i*_ obtained using specific realizations of random connectivity in Eq. (9), the rank-one approximation provides predictions for the activity of single units in *individual instances* of locally-defined networks. Moreover, the rank-one theory predicts that both the population-average and the standard deviation of activations in the network are proportional to *κ*.

The values taken by the latent variable *κ* can be determined by projecting Eq. (13) onto **m** and inserting Eq. (15) (Methods Secs. 2.6.1, 2.6.2). This leads to a closed equation for the dynamics of *κ*(*t*):

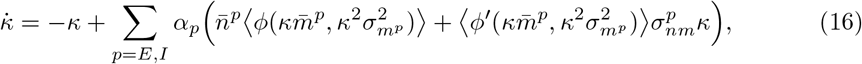

where the brackets denote a Gaussian average (see Eq. (110)). The steady state then obeys

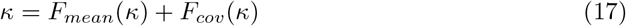

where

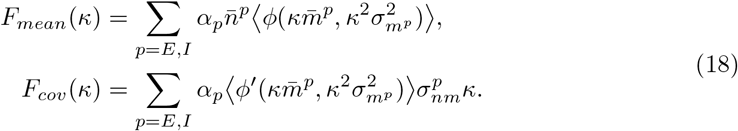

The two terms in the r. h. s. of Eq. (17) show that the contributions of recurrent synaptic inputs to the latent dynamics *κ* come from two sources: (i) the population means of the left and right connectivity eigenvectors 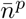 and 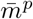 that contribute to *F*_*mean*_(*κ*) (Eqs. (52)-(54)); (ii) the covariance 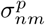 between the left and right connectivity eigenvectors that contributes only to *F*_*cov*_(*κ*). In the low-rank approximation of the locally-defined E-I connectivity, these two terms have distinct origins: the mean comes from the independent components of the connectivity (Eqs. (53), (54)); while the covariance comes from reciprocal correlations between connections (Eqs. (94), (95)). We therefore next examine separately the effects on dynamics of these two connectivity components.

#### 1.5.1 Independently generated local connectivity

When synaptic connections are generated independently from a Gaussian distributions based on the identities of pre- and post-synaptic populations, the rank-one approximation of connectivity leads to uncorrelated left and right connectivity vectors **n** and **m**, so that 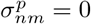 for *p* = *E, I*. In consequence, only the first term is present in the r. h. s. of Eq. (17), and the fixed point of the latent dynamics is given by a difference between excitatory and an inhibitory feedback (Eq. (114)):

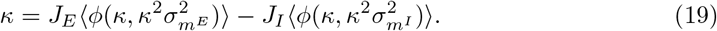

As long as the mean inhibition *J*_*I*_ is strong enough to balance the mean excitation *J*_*E*_, Eq. (19) predicts a single fixed point. As *J*_*E*_ is increased, positive feedback begins to dominate and leads to a bifurcation to a bistable regime for the latent dynamic variable *κ* (Figs 5A-C, S2 Appendix).

**Fig 5.**
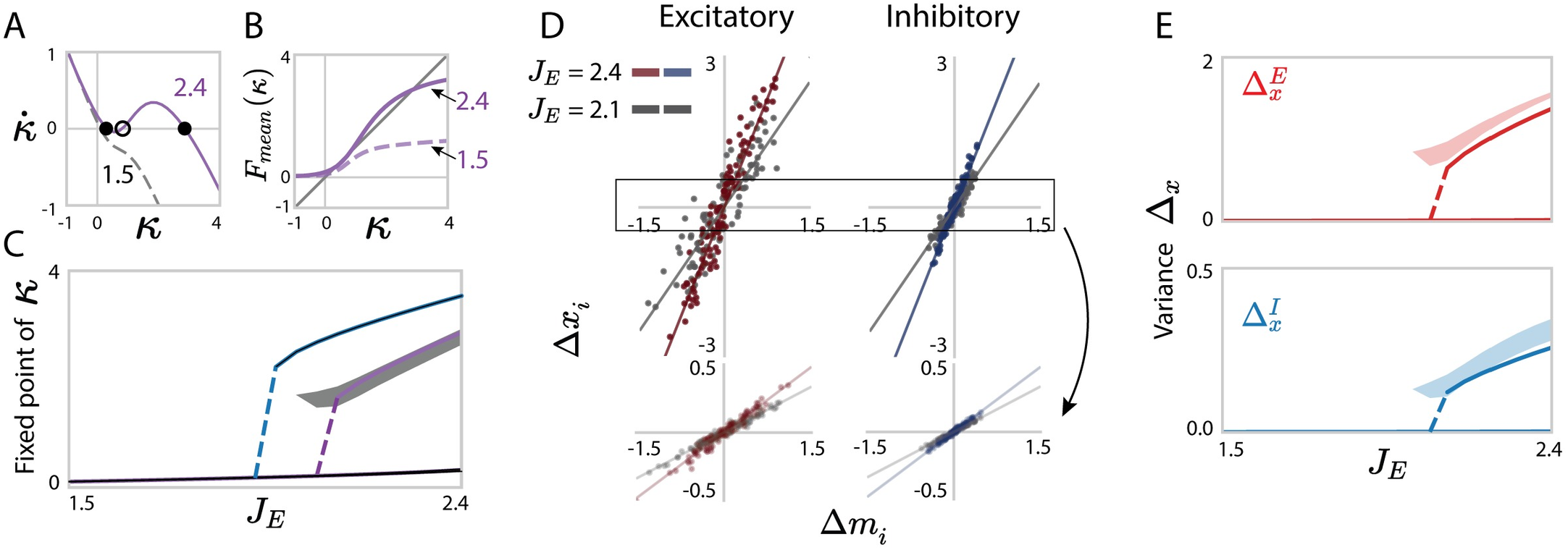
Predicting low-dimensional dynamics using a rank-one approximation of networks with independent Gaussian connectivity. (A) Fixed points of the latent variable *κ* in the rank-one approximation. The lines show the dynamics 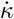 as function of *κ*, predicted by Eq. (16) (solid line: *J*_*E*_ = 2.4; dashed line: *J*_*E*_ = 1.5). The intersections with *y* = 0 correspond to fixed points (filled dots: stable; unfilled dot: unstable). (B) Contribution of mean connectivity to the latent dynamics, *F*_*mean*_(*κ*) in Eq. (18), for two values of *J*_*E*_. (C) Bifurcation diagram for increasing *J*_*E*_: analytical predictions of Eq. (19) compared with simulations of the full network with locally-defined connectivity. Blue line: analytical prediction including only the mean part of the connectivity (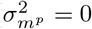 in Eq. (19)); purple line: analytical prediction including the first-order perturbation term in the rank-one approximation; gray: projection of simulated activity **x** onto the connectivity vector **m** computed by perturbation theory (Eq. (102)). (D) Comparison between predictions and simulations for the activity of individual units in a given realization of the random connectivity. For each unit *i*, a dot shows the deviation Δ*x*_*i*_ of its steady-state activity from the population average, against its value Δ*m*_*i*_ of the perturbed part of the connectivity vector **m** (Eq. (15)). The low-rank theory predicts Δ*x*_*i*_ = *κ*Δ*m*_*i*_. Red and blue scatters show excitatory or inhibitory populations, each for several values of *J*_*E*_. Lines represent *y* = *κx*, where *κ* is obtained from Eq. (19). Upper panels show the result in a realization with a high fixed point, bottom panels show the result in a realization with a low fixed point. (E) Comparison between the predictions (solid lines) and simulations (shaded areas) for the population-averaged variances of Δ*x*_*i*_. Shaded areas show mean±std. Network parameters: *N*_*E*_ = 4*N*_*I*_ = 1200, *J*_*I*_ = 0.6, *g*_*EE*_ : *g*_*EI*_ : *g*_*IE*_ : *g*_*II*_ = 1.0 : 0.5 : 0.2 : 0.8 and *g*_*EE*_ = 0.8. The transfer function *ϕ* has parameter *θ* = 1.5.

This bistability due to positive feedback is expected on the basis of mean connectivity alone. Indeed replacing the connectivity matrix by its mean 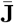 is equivalent to a rank-one approximation with 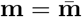 and 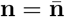 which lead to Eq. (113) with 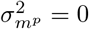 for *p* = *E, I*. The additional first-order perturbation term in the rank-one approximation (Eqs. (8), (9)) additionally takes into account fluctuations in the connectivity, which leads to a non-zero 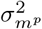, and modifies the fixed points predicted by
Eq. (19). In consequence, the bifurcation to bistability takes place at higher values of *J*_*E*_ than predicted from mean connectivity alone (purple lines compared to blue lines in Fig 5C).

More importantly, we find that the first-order perturbation in the rank-one approximation accurately predicts the firing rates of individual neurons for specific instances of the random, locally-defined connectivity (Fig 5D), and therefore the variance of the steady state of population dynamics 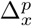. In particular, cell-type dependent variances in the synaptic connectivity, lead to distinct variances 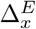 and 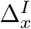 for excitatory and inhibitory populations (Eqs. (88), (129), Fig 5E).

Note that the independently generated local connectivity can be treated analytically without resorting to a rank-one approximation, by using a different variant of mean-field theory originally developed for randomly connected networks. [30, 52]. That theory is not perturbative, and takes into account an additional term in the variance (see Methods Sec. 2.6.3 and S3 Appendix for more details). However, in contrast to the rank-one approximation, it does not predict the activity of individual neurons, and is challenging to extend beyond independent random connectivity.

#### 1.5.2 Reciprocal motifs

We next turn to the predictions of the rank-one approximation for dynamics resulting from locally-defined connectivity with reciprocal motifs. In this case, the additional reciprocal correlations in the random part of the connectivity lead to a non-zero covariance *σ*_*nm*_ between the connectivity vectors **n** and **m** in the rank-one approximation (Eqs. (90), (95)). This covariance in turn generates an additional feedback component in the dynamics of the latent variable, the second term in the r. h. s. of Eq. (17).

Specifically, positive reciprocal correlations combined with excitation-dominated connectivity enhance positive feedback with respect to mean connectivity alone. As a result, progressively increasing the reciprocal correlations can therefore induce a bifurcation to bistability, even if the mean excitation is not sufficient by itself to support two stable states (Figs 6A-C). This is a major novel effect of reciprocal motifs on collective dynamics. As in the case of independent connectivity, we moreover found that the perturbative term in the rank-one approximation correctly predicts the activity of individual neurons in specific realizations of the connectivity (Fig 6D), and therefore also the cell-type dependent variances of activity (Fig 6E).

**Fig 6.**
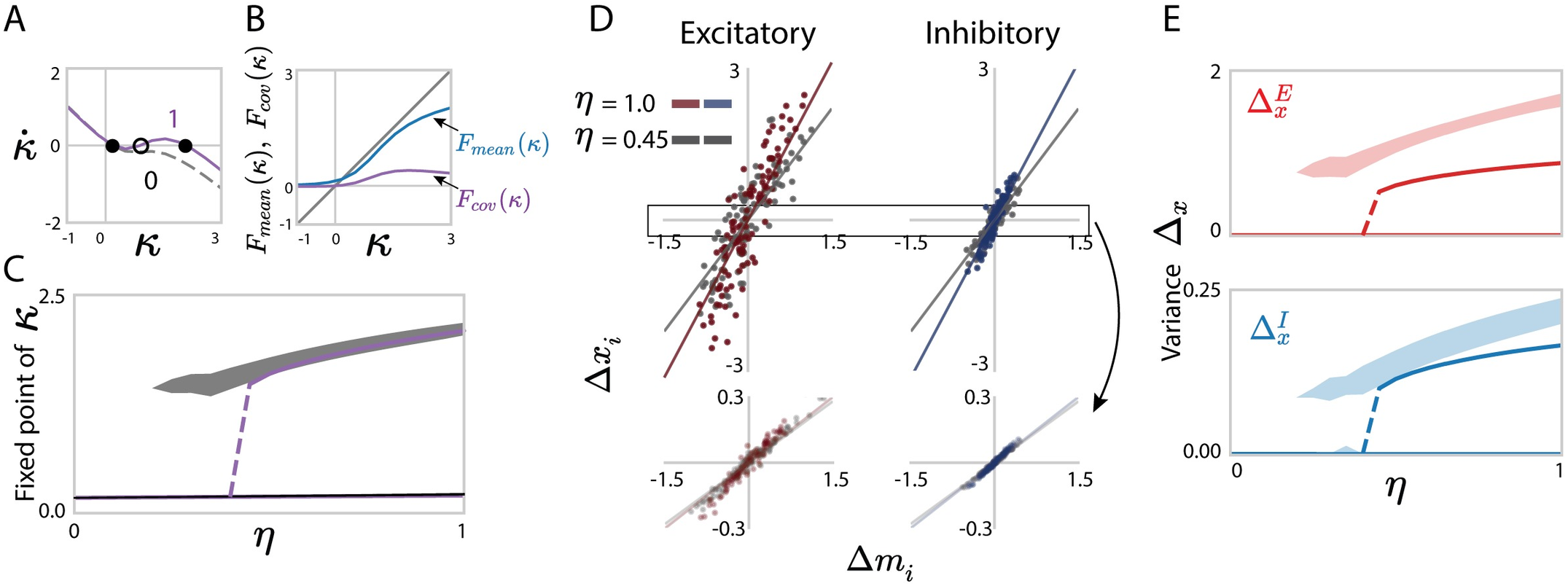
Predicting low-dimensional dynamics using a rank-one approximation of networks with homogeneous reciprocal motifs. (A) Influence of reciprocal correlations on fixed points of the latent variable *κ* in the rank-one approximation. The lines show the dynamics 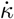 as function of *κ*, predicted by Eq. (16) (solid line: *η* = 1; dashed line: *η* = 0). The intersections with *y* = 0 correspond to fixed points (filled dots: stable; unfilled dot: unstable). (B) Comparison of the contributions of mean connectivity *F*_*mean*_(*κ*) and covariance *F*_*cov*_(*κ*) to the latent dynamics of *κ* (Eq. (18)) for *η* = 1. (C) Bifurcation diagram for increasing *η* at fixed *J*_*E*_. Solid purple lines: analytical predictions of Eqs. (17), (18); gray areas: projection of simulated activity **x** onto the connectivity vector **m** computed by perturbation theory (Eq. (102)). (D) Comparison between predictions and simulations for the activity of individual units in a given realization of the random connectivity. For each unit *i*, a dot shows the deviation Δ*x*_*i*_ of its steady-state activity from the population average, against its value Δ*m*_*i*_ of the perturbed part of the connectivity vector **m** (Eq. (15)). The low-rank theory predicts Δ*x*_*i*_ = *κ*Δ*m*_*i*_. Red and blue scatters show excitatory or inhibitory populations, each for two values of *η*. Lines represent *y* = *κx*, where *κ* is obtained from Eqs. (17), (18). Upper panels show the result in a realization with a high fixed point, bottom panels show the result in a realization with a low fixed point. (E) Comparison between the predictions (solid lines, Eq. (133)) and simulations (shaded areas) for the population-averaged variances of Δ*x*_*i*_. Shaded areas show mean±std. Network parameters: *N*_*E*_ = 4*N*_*I*_ = 1200, *J*_*E*_ = 1.9, *J*_*I*_ = 0.6, *g*_*EE*_ : *g*_*EI*_ : *g*_*IE*_ : *g*_*II*_ = 1.0 : 0.5 : 0.2 : 0.8 and *g*_*EE*_ = 0.8. The transfer function *ϕ* has parameter *θ* = 1.5.

More generally, our rank-one approximation allows us to describe the latent dynamics when the degree of reciprocal correlation depends across the pre- and post-synaptic populations (Results Sec. 1.4). Such heterogeneity in reciprocal correlations can enhance different types of feedback. For example, antisymmetric connectivity within inhibitory populations (*η*_*II*_ *<* 0) disinhibits excitatory population and thus facilitates bistable transitions (Figs 7A-C) compared to networks with homogeneous reciprocal correlations. In contrast, excitation-dominated connectivity with homogeneous negative reciprocity (*η <* 0) generate negative feedback and therefore suppress the global dynamics from bistable state to quiescent (Figs 7D-F).

**Fig 7.**
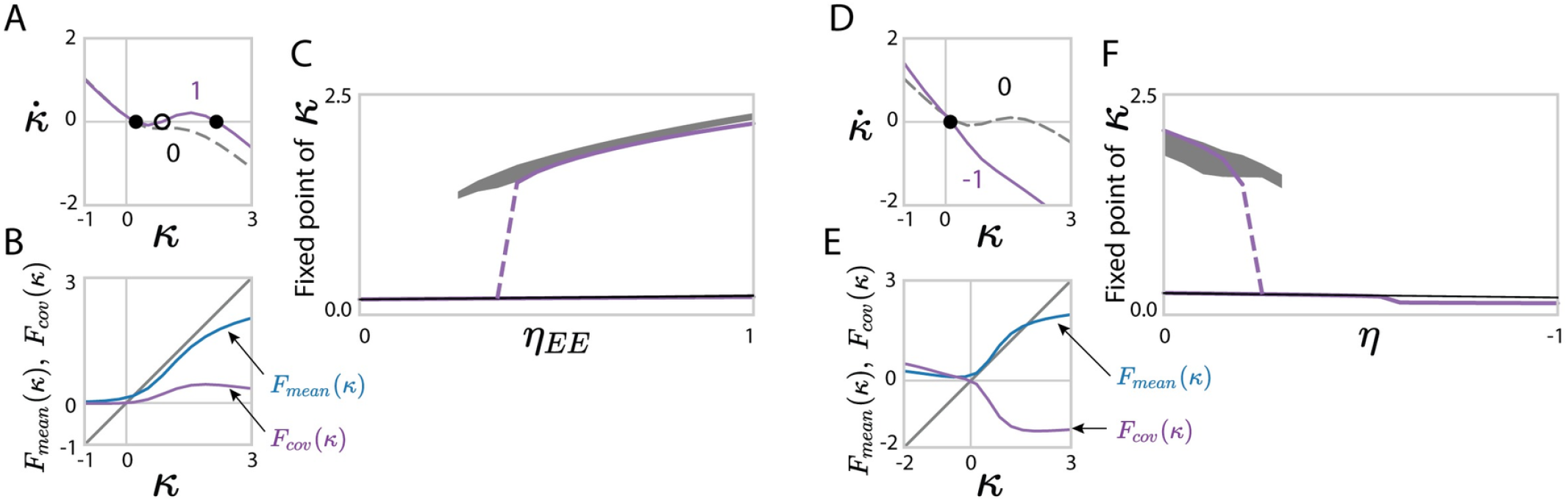
Predictions for low-dimensional dynamics using a rank-one approximation of networks with non-homogeneous and anti-symmetric reciprocal motifs. (A-D) Fixed points of latent dynamics in networks with heterogeneous, cell-type-dependent reciprocal correlations: *η*_*EE*_ = *η*_*EI*_ = −*η*_*II*_. Network parameters: *N*_*E*_ = 4*N*_*I*_ = 1200, *g*_*EE*_ : *g*_*EI*_ : *g*_*IE*_ : *g*_*II*_ = 1.0 : 0.5 : 0.2 : 0.8, *g*_*EE*_ = 0.8, *J*_*E*_ = 1.9 and *J*_*I*_ = 0.6. (E-H) Fixed points of latent dynamics in networks with homogeneously anti-symmetric motifs, *η*_*pq*_ = *η* ∈ [−1, 0]. Network parameters: *N*_*E*_ = 4*N*_*I*_ = 1200, *g*_*EE*_ : *g*_*EI*_ : *g*_*IE*_ : *g*_*II*_ = 1.0 : 0.5 : 0.2 : 0.8, *g*_*EE*_ = 0.8, *J*_*E*_ = 2.2 and *J*_*I*_ = 0.6. In C and F, gray areas show projections of simulated activity **x** onto the connectivity vector **m** computed by perturbation theory (Eq. (102)), shaded areas show mean±std.

Importantly, describing the role of reciprocal correlations on latent dynamics relies on our global low-rank approximation of locally-defined connectivity. In particular, the effects of such correlations cannot be captured by considering only mean connectivity and population-averaged activity (first term in the r. h. s. of Eq. (17)). Moreover, including reciprocal correlations in classical mean-field approaches to randomly connected networks is technically challenging [18].

### 1.6 Extension: E-I networks with sparse connectivity

In previous sections, we examined locally-defined connectivity generated using Gaussian distributions of individual synaptic weights (function *f*^*pq*^ in Eqs. (1)). Our results for the low-rank approximation of locally-defined connectivity are however independent of the precise form of the distribution *f*^*pq*^. In particular, our finding that the resulting low-rank structure obeys Gaussian-mixture statistics is universal, in the sense that it is valid for any distribution *f*^*pq*^ for which the central limit theorem holds (see Discussion). To illustrate this universality, here we turn to networks with sparse connectivity, generated from Bernoulli distributions *f*^*pq*^ taking values 0 and *A*_*q*_ (where *A*_*q*_ = *A*_*E*_, *A*_*I*_ refer to strengths of excitatory and inhibitory connections), with a uniform fraction *c* of non-zero connections (see Eq. (26)). The statistics of the corresponding rank-one approximation are fully determined by the mean, variance and covariance of synaptic weights in the excitatory and inhibitory populations. Here we compare the predictions of our perturbative approximation with direct simulations of full-rank networks with locally-defined connectivity. We first consider independently generated connectivity, and then turn to reciprocal motifs.

#### 1.6.1 Independently generated sparse connectivity

For networks with independently generated sparse connectivity, the mean and variance of individual synaptic weights are given by *cA*_*q*_ and 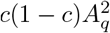, where *q* = *E, I* refers to the population of the presynaptic neuron. Previous works have shown that such sparse networks have a random, circularly distributed bulk of random eigenvalues with a radius *r*_*g*_ determined by the overall variance of synaptic weights (Eq. (148), Fig 8A) [53, 54], as expected from the universality theorem for random matrices [49]. The overall mean of the synaptic weights instead determines the mean outlying eigenvalue (Fig 8A). In sparse networks, the main novelty with respect to the Gaussian case is that the mean and variance of synaptic weights are not independent parameters, but are instead both set by the synaptic couplings *A*_*E*_ and *A*_*I*_ as well as the network’s sparsity *c*. In consequence, varying these couplings changes both the radius of the bulk and the outlying eigenvalue, and can lead to intersections where the outliers dip into the bulk (see S4 Appendix for details).

**Fig 8.**
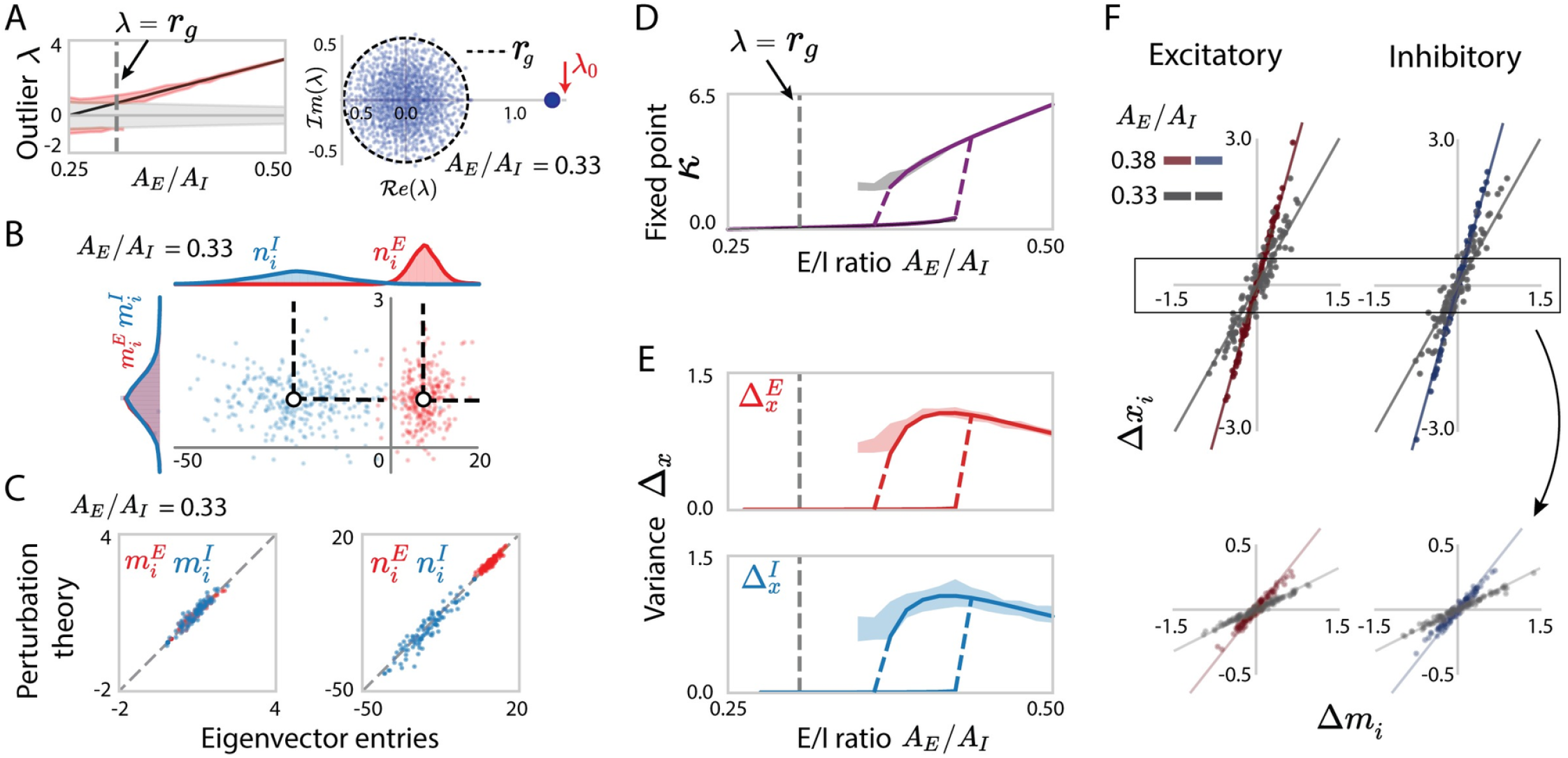
Rank-one approximation and predicted low-dimensional dynamics for sparse excitatory-inhibitory networks. (A) Left: Comparison of the predicted eigenvalue outlier *λ*_0_ = *c*(*N*_*E*_*A*_*E*_ + *N*_*I*_*A*_*I*_) (black line) with finite-size simulations (red area shows mean±std for 30 realizations). The gray area represents the area covered by the eigenvalue bulk. Right: example spectrum of one realization of connectivity matrix with E/I ratio *A*_*E*_*/A*_*I*_ = 0.33, and the radius *r*_*g*_ of the eigenvalue bulk computed from the statistically equivalent Gaussian connectivity (see S4 Appendix). (B) Scatter plot showing for each neuron *i* its entry *n*_*i*_ on the left eigenvector against its entry *m*_*i*_ on the right eigenvector. Red and blue colours represent respectively excitatory and inhibitory neurons. The white dots and the dash lines respectively indicate the means and covariances for each population obtained from simulations. For visualization purposes, the *x*−and *y*-axis are scaled unequally. (C) Comparison between eigenvector entries obtained from direct eigen-decomposition of **J** with predictions of perturbation theory (Eqs. (8), (9)). (D) Bifurcation diagram for increasing the ratio *A*_*E*_*/A*_*I*_ : analytical predictions of Eq. (115) compared with simulations of the full network with locally generated sparse connectivity. Purple line: analytical prediction including the first-order perturbation term in the rank-one approximation; gray: projection of simulated activity onto the connectivity vector **m** computed by perturbation theory Eq. (102). (E) Comparison between the predictions (solid lines) and simulations (shaded areas) for the population-averaged variances of Δ*x*_*i*_. Shaded areas show mean std. ±(F) Comparison between predictions and simulations for the activity of individual units in a given realization of the sparse connectivity. For each unit *i*, a dot shows the deviation Δ*x*_*i*_ of the steady-state activity from the population average, against the corresponding value Δ*m*_*i*_ of the perturbed part of the connectivity vector **m** (Eq. (15)). The low-rank theory predicts Δ*x*_*i*_ = *κ*Δ*m*_*i*_. Red and blue scatters show excitatory or inhibitory populations, each for two values of the ratio *A*_*E*_*/A*_*I*_. Lines represent *y* = *κx*, where *κ* is obtained from Eqs. (17), (18). Upper panels show the result in a realization with a high fixed point, bottom panels show the result in a realization with a low fixed point. The gray vertical dashed line in A left, D, E correspond to the critical point *c* at which the absolute value of the outlier is equal to the radius of the eigenvalue bulk. Network parameters: *N*_*E*_ = 4*N*_*I*_ = 800, *c* = 0.3, *A*_*E*_ = 0.025. The transfer function *ϕ* has parameter *θ* = 1.5.

As long as the outlier lies outside of the eigenvalue bulk, our perturbative approximation Eq. (67) predicts well the individual entries of the right- and left-eigenvectors corresponding to the outlier from the fluctuation matrix **Z** (Fig 8C). As expected, the resulting statistics of eigenvectors entries are well described by a Gaussian-mixture distribution with parameters fully determined by the mean and variance of the synaptic weights (Fig 8B).

Our predictions for the low-dimensional dynamics based on the rank-one approximation therefore directly extend to sparse networks. Comparing with direct simulations, we found that Eq. (115) predicts well the global latent variable *κ* obtained by projecting the activity **x** onto the approximated rank-one eigenvector **m** (Eq. (102)). As the E/I ratio is increased, positive feedback increases, and the latent variable *κ* undergoes a transition from a single fixed point to two bistable states (Fig 8D).

For individual realizations of the sparse connectivity, the rank-one approximation **x** = *κ***m** predicts well the activation *x*_*i*_ of individual neurons in the simulations (Fig 8F). From the statistics of the right eigenvector **m** (Eq. (89)), our analysis moreover predicts the heterogeneity of activity in terms of population-averaged variances 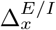 (Fig 8E). This heterogeneity is identical in excitatory and inhibitory populations, as their right eigenvectors have identical fluctuations (Figs 8C, E).

#### 1.6.2 Sparse EI networks with reciprocal motifs

For sparse E-I networks, we generate reciprocal motifs by introducing a fraction *ρ*_*pq*_ of reciprocally connected pairs of neurons. Together with the sparsity *c* and the synaptic strengths, the parameter *ρ*_*pq*_ determines the cell-type dependent reciprocal correlation *η*_*pq*_ (Methods Sec. 2.1.2 Eqs. (32), (37), Fig 9A).

**Fig 9.**
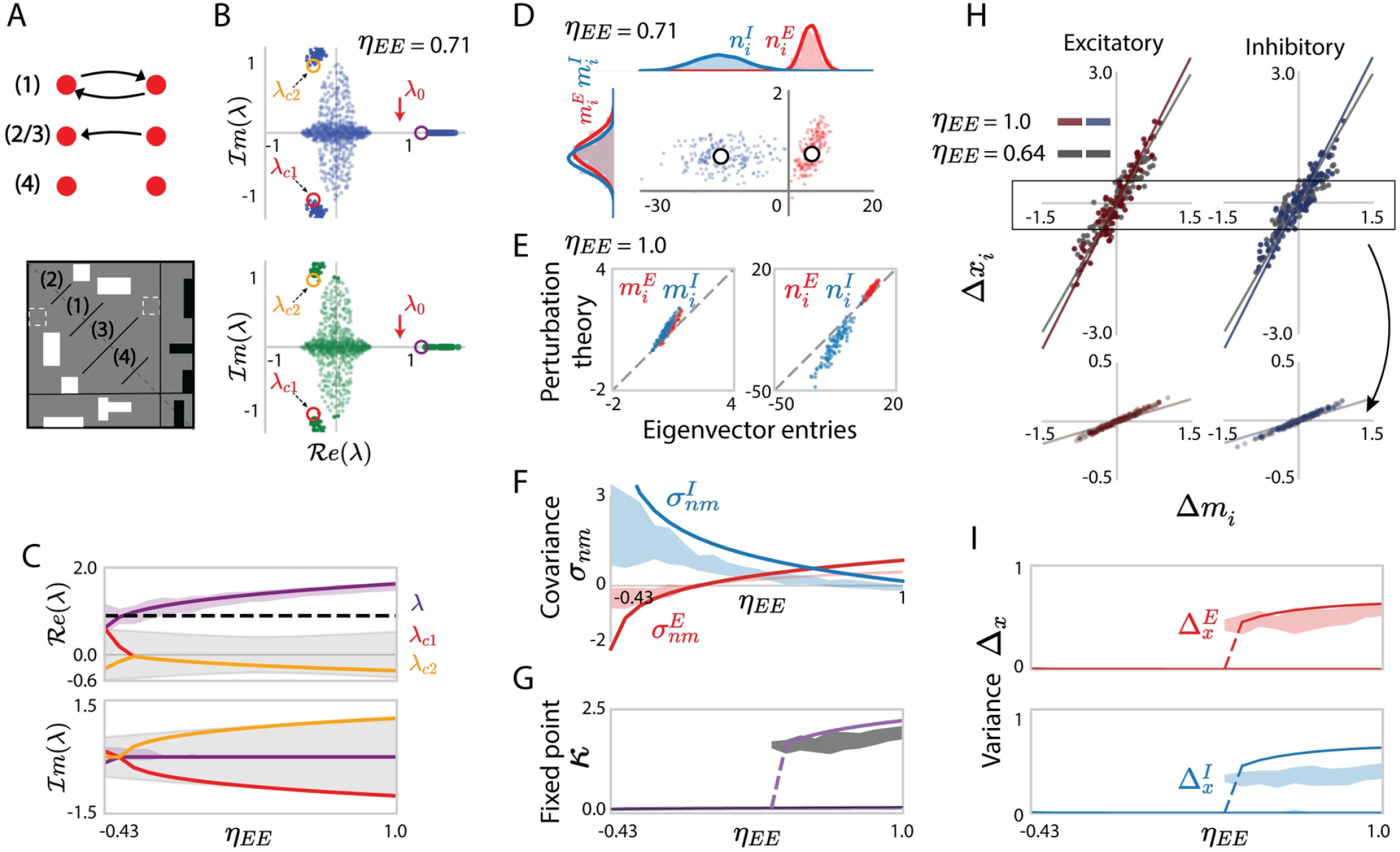
Characterizing connectivity statistical properties and low-dimensional dynamics for the sparse network with reciprocal motifs. (A) Schematics of a sparse EI network with four forms of paired connections. White and black rectangles represent the non-zero excitatory and inhibitory sparse connections. (B) Eigenvalue spectrum of the sparse connectivity (upper panel) and from the equivalent Gaussian connectivity (bottom panel) with reciprocal motifs. Cell-type-dependent reciprocal correlations are *η*_*EE*_ =−0.71, *η*_*EI*_ = −0.71, *η*_*II*_ = −0.43 in both connectivity matrices, continuous eigenvalue bulks show eigenvalues for one realization of the network connectivity. Red arrows point to the unperturbed eigenvalue *λ*_0_. Outlying eigenvalues are shown for 30 realizations of the network connectivity. Coloured circles are the eigenvalues predicted using determinant lemma. (C) Comparison of the eigenvalues from the finite-size simulation of the sparse connectivity, with the predictions of the determinant lemma as progressively increasing the reciprocal correlation *η*_*EE*_ (−*η*_*EI*_). The coloured solid lines show the roots of the third-order polynomial in Eq. (7). The purple area indicates the empirical distribution of the dominant outlier, while the black dashed line is the eigenvalue *λ*_0_ of the corresponding independent sparse connectivity matrix (Eq. (76)). The gray areas correspond to the areas covered by the eigenvalue bulk. (D) Scatter plot showing for each neuron *i* its entry *n*_*i*_ on the left eigenvector against its entry *m*_*i*_ on the right eigenvector. Red and blue colours represent respectively excitatory and inhibitory neurons. The white dots indicate the means for each population obtained from simulations. For visualization purposes, the *x*- and *y*-axis are scaled unequally. (E) Comparison between eigenvector entries obtained from direct eigen-decomposition of **J** with predictions of perturbation theory (Eqs. (8), (9)). (F) Comparison between the population covariance 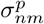 of the entries on the left and right connectivity eigenvectors to different populations (coloured areas) and the predictions of perturbation theory (coloured lines, Eq. (97)). (G) Bifurcation diagram for increasing the reciprocal correlation *η*_*EE*_ (−*η*_*EI*_): analytical predictions of Eq. (117) compared with simulations of the full network with locally generated sparse connectivity and reciprocal motifs. Purple line: analytical prediction including the first-order perturbation term in the rank-one approximation; gray: projection of simulated activity onto the connectivity vector **m** computed by perturbation theory Eq. (102). (H) Comparison between predictions and simulations for the activity of individual units in a given realization of the sparse connectivity. For each unit *i*, a dot show the deviation Δ*x*_*i*_ of the steady-state activity from the population average, against its value Δ*m*_*i*_ of the perturbed part of the connectivity vector **m** (Eq. (15)). The low-rank theory predicts Δ*x*_*i*_ = *κ*Δ*m*_*i*_. Red and blue scatters show excitatory or inhibitory populations, eachfor two values of the reciprocal correlation *η*_*EE*_. Lines represent *y* = *κx*, where *κ* is obtained from Eqs. (17)-(18). Upper panels show the result in a realization with a high fixed point, bottom panels show the result in a realization with a low fixed point. (I) Comparison between the predictions (solid lines) and simulations (shaded areas) from the population-averaged variances of Δ*x*_*i*_, shaded areas show mean±std. In C, F, G, I, the reciprocal correlations *η*_*EE*_ = −*η*_*EI*_ progressively increase from −0.43 to 1.0 while keeping *η*_*II*_ = −0.43 constant (*ρ*_*EE*_ = *ρ*_*EI*_ increase from 0 to 1 and *ρ*_*II*_ = 0 is fixed). Network parameters: *N*_*E*_ = 4*N*_*I*_ = 800, *c* = 0.3, *A*_*E*_ = 0.023, *A*_*E*_*/A*_*I*_ = 0.3. The transfer function *ϕ* has parameter *θ* = 1.5.

We first examine the effect of the reciprocal motifs on the statistical properties of the eigenvalues and eigenvectors. The spectrum still consists of continuous eigenvalues and a discrete outlier. Since in the independent case the outlier depends only on *A*_*E/I*_ and sparsity *c*, here we fix these two variables but increase *η*_*EE*_ = −*η*_*EI*_ while keeping *η*_*II*_ constant. As in the case of dense, Gaussian networks, the outlier increases with the increasing reciprocal correlation and deviates from the outlier of the corresponding independent sparse connectivity matrix (Figs 9B, C). Moreover, in the large network limit, we find that if means, variances, and reciprocal correlations are identical, dense Gaussian connectivity leads to the same eigenvalue spectrum as the sparse connectivity (Methods Secs. 2.1.2, 2.1.3 Eqs. (36), (37)). We furthermore mathematically predict two additional conjugate eigenvalue outliers generated by the reciprocal connections in the sparse case (Eqs. (80), (86), (87), Fig 9B).

As for uncorrelated connectivity, perturbation theory predicts the individual left and right eigenvector entries, which altogether follow Gaussian statistics as expected (Figs 9D, E). Importantly, reciprocal correlations induce a non-zero covariance *σ*_*nm*_ between the entries *m*_*i*_ and *n*_*i*_ of the right and left eigenvectors (Eq. (97), Fig 9F).

Finally, we examine the population dynamics in the sparse network with reciprocal motifs using the low-rank approximation derived above. The reciprocal motifs in the example network generate an overall positive feedback. Therefore, gradually increasing the reciprocal correlation *η*_*EE*_ (−*η*_*EI*_) in the example network induces a bifurcation into bistability (Eq. (117), Fig 9G). Analogous to the predictions of individual eigenvector entries, the low-rank approximation gives analytical predictions for the activity of individual neurons in specific connectivity realizations (Fig 9H), and hence the cell-type dependent variances of neuronal activation 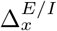 (Fig 9I) obtained from finite-size simulations of the original sparse networks.

## Discussion

In this work, we unified two different descriptions of connectivity in multi-population networks and thereby connected two broad classes of models. Starting from local statistics of synaptic weights, we approximated the resulting connectivity matrix in terms of a low-rank structure. The obtained, approximate low-rank network model then allowed us to determine the influence of the local connectivity motifs on the global low-dimensional dynamics.

A key ingredient in our approach is a low-rank approximation of the locally-defined connectivity matrix. Approximating an arbitrary full-rank matrix by a rank-*R* one is a classical problem in numerical analysis, for which a number of different methods are available depending on the objective of approximation [55]. The most common method is to perform a singular value decomposition (SVD), and keep the top *R* terms [56]. This method minimizes the Frobenious norm of the difference between the original matrix and its low-rank approximation. Our goal in this study was however to obtain a low-rank approximation that preserves the dominant eigenvalues of the original matrix, as these eigenvalues determine the autonomous dynamics in the network. An SVD-based approximation preserves the top singular values, but in general not the top eigenvalues. We therefore opted for an approximation based on truncated eigen-decomposition. When studying input-driven and transient dynamics, different methods for low-rank approximation may be more appropriate, and are a topic of active research [57–61]

To perform the eigen-decomposition of excitatory-inhibitory connectivity matrices, we leveraged the fact that they can be expressed as a sum of a block-like deterministic low-rank matrix and a full-rank random matrix with zero-mean [40]. The eigen-spectrum of such matrices in general consists of a continuously-distributed bulk and discrete outliers [44, 45]. While a number of works have examined the bulk of the eigenvalue spectrum for random matrices [11, 18, 32, 44, 49, 52, 62, 63], the outliers, and in particular the corresponding eigenvectors have to our knowledge received less attention. The main technical novelty in this work is the use of matrix perturbation theory [47, 64] to approximate the eigenvectors corresponding to the outliers in the eigenspectrum of the locally-defined connectivity matrices. A key output of this approach is the finding that entries of the left- and right-eigenvectors follow multivariate Gaussian distributions, the statistics of which depend on the population the neurons belong to. This result provides a general theoretical mapping from locally-defined multi-population models to Gaussian-mixture low-rank networks [31, 33]. It however holds only as long as the distribution of synaptic weights satisfies the assumptions of the central limit theorem. This in particular excludes heavy tailed distributions often found in experimental studies [1, 65].

In the networks we considered, the non-random structure in connectivity comes only from the multi-population organization. More specifically, the low-rank skeleton of the locally-defined connectivity matrix is fully specified by the mean synaptic weights between different populations (Eq. (3)). This mean connectivity structure largely controls the outlying eigenvalue, and the average values of the corresponding eigenvector entries. The random part of the connectivity and reciprocal motifs can modify the outlying eigenvalue, and add heterogeneity as well as correlations to this underlying structure. Our perturbative theory allows us to quantify these effects and predict dynamics on a single-neuron basis. This approach can be directly extended to networks with additional structure, in which the low-rank skeleton is not solely determined by the mean connectivity but possibly by more general patterns.

A key insight from our study is a general relationship between reciprocal motifs in locally-defined connectivity and overlaps among connectivity vectors in low-rank networks. Indeed, we have shown that correlations between reciprocal synaptic weights generate overlaps beyond the mean in the corresponding low-rank approximation (Eq. (93)). Conversely, zero-mean overlaps between connectivity vectors in a low-rank model necessarily imply non-vanishing reciprocal correlations (Eq. (41) and S5 Appendix). Since overlaps between connectivity vectors determine the autonomous recurrent dynamics in low-rank networks, this relationship allowed us to quantify how reciprocal connectivity motifs contribute to network dynamics.

Local statistics of synaptic connectivity are believed to play an important role in the global network dynamics [1, 17, 32, 62]. Our study, provides a mathematical theory that relates the local connectivity statistics to global recurrent dynamics through a low-rank approximation. This computational framework is not restricted to the reciprocal motifs that we have emphasized in this work, but can be extended to various forms of local connectivity motifs [10]. As a result, our framework can be applied more broadly to study experimentally-obtained connectivity databases and connectivity consisting of different types of motifs. [6, 7, 15].

## 2 Materials and methods

Throughout this study, we consider recurrent networks of *N* neurons and denote by **J** the recurrent connectivity matrix, where *J*_*ij*_ is the synaptic strength of the connection from neuron *j* to neuron *i*.

### 2.1 Locally-defined multi-population connectivity

In this section, we introduce a first class of connectivity models, in which the synaptic couplings are generated based on *local* statistics determined by the identity of pre- and post-synaptic neurons. The *N* neurons in the network are organized in *P* populations, where population *p* has *N*_*p*_ neurons. Denoting by *p* and *q* the populations neurons *i* and *j* belong to, the value of the synaptic coupling *J*_*ij*_ is drawn randomly from a distribution in which statistics depend on the pre- and postsynaptic population *q* and *p*. The full connectivity matrix **J** therefore has a block structure, in the sense that all connections within the same block share *identical* statistics.

We examine two variants of this model class: (1) ***independent*** random connectivity [52]; (2) connectivity with ***reciprocal motifs*** [10, 66]. In each case, we examine two specific examples of distributions of synaptic strengths, *Gaussian*, and *Sparse* distributions.

#### 2.1.1 Independent random connectivity

For networks with independent random connectivity, the recurrent connections *J*_*ij*_ are sampled independently for each (*i, j*) pair from

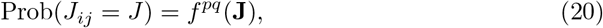

where *f*^*pq*^ denotes a probability density function, and *q, p* are the pre- and post-synaptic populations. Separating the mean and random components, for an arbitrary distribution Eq. (20) can be re-expressed as

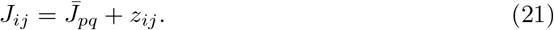

Here 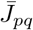 is the mean value of the connections from population *q* to population *p*, and *z*_*ij*_ is the remaining zero-mean random part of each connection. Defining 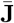 as the *N* × *N* deterministic matrix consisting of mean values, and **Z** as the noise matrix consisting of the random parts *z*_*ij*_, the connectivity matrix **J** can be written as

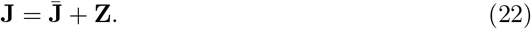

The matrix 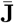 is of size *N*×*N* and consists of *P* ^2^ blocks with identical values within each block. The rank of 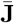 is therefore at most *P* [40]. In contrast, the noise matrix **Z** is in general of rank *N*. The full connectivity matrix **J** can then be interpreted as a rank-*P* deterministic matrix perturbed by the random matrix **Z** with block-dependent statistics.

In the case of Gaussian connectivity, connections from population *q* to population *p* are sampled independently from a Gaussian distribution.

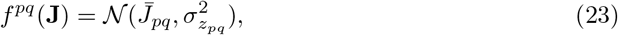

with variances

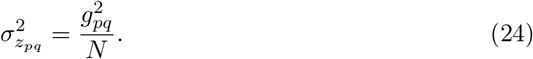

The noise matrix **Z** therefore has *block-structured variances* 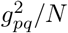 that we specify by a *P* × *P* matrix **G**_*m*_:

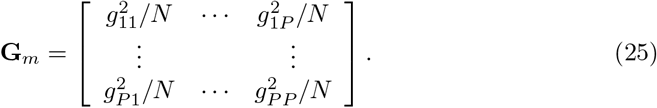

In the case of sparse connectivity, 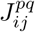 is a Bernoulli random variable. The connectivity weights 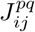 from population *q* to population *p* are non-zero with probability *c*_*pq*_ and zero otherwise. All non-zero connection weights within a block take the same value *A*_*pq*_, so that analogously to Eq. (20), the sparse connectivity is defined as

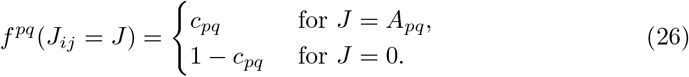

The mean connectivity weight between populations *p, q* is then

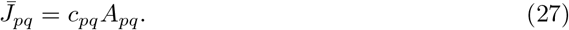

and the variance of the remaining random part *z*_*ij*_ is

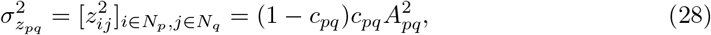

to simplify the parameters in sparse networks, we assume that *A*_*pq*_ depend only on presynaptic population *q*, and that the connection probability *c*_*pq*_ is a homogeneous network parameter independent of *p, q* that we denote by *c*.

#### 2.1.2 Reciprocal connectivity motifs

To go beyond independent connectivity, we consider pairwise motifs, i. e. correlations between reciprocal pairs of weights *J*_*ij*_ and *J*_*ji*_. We quantify this correlation using the normalized covariance *η*_*ij*_ defined as.

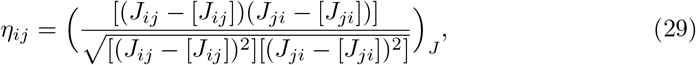

where [·] denotes the average over the full connectivity distribution. Reciprocal connections are fully independent when *η*_*ij*_ = 0 for all *i, j*, fully symmetric when *η*_*ij*_ = 1 and fully anti-symmetric when *η*_*ij*_ =−1.

Our key assumption is that the statistics of connectivity are block-like, implying that all pairs of connections between populations *p, q* share the same correlation coefficient *η*_*pq*_, so that the statistics are defined by a *P* × *P* reciprocal correlation matrix ***η***_*m*_

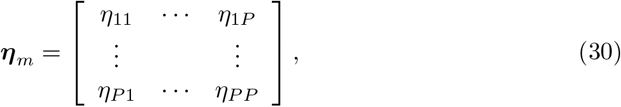

where, by definition *η*_*pq*_ = *η*_*qp*_.

For Gaussian statistics, we generate connectivity matrices with a specified set of *η*_*pq*_ in the following manner. We first generate an *N*×*N* matrix **Y**′ with entries independently sampled from the normal distribution𝒩 (0, 1). We then form a linear combination **Y**′ and its transpose **Y**′^⊤^ to generate a matrix **Y** with reciprocal correlations *η*_*pq*_. Specifically, we set

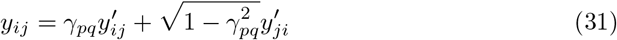

with 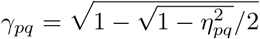 for *η*_*pq*_ *>* 0, and 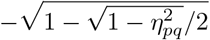 for *η*_*pq*_ *<* 0. Finally, we scale each block by 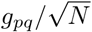 to obtain the random connectivity component **Z**, which is added to the mean connectivity component 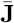 to finally obtain the full connectivity matrix **J**.

For *sparse* networks, we first generate a connectivity matrix without reciprocal correlations. We then consider the upper triangle of this matrix, randomly select a fraction *ρ*_*pq*_ of the non-zero connections *J*_*ij*_ with value *A*_*q*_ and set their reciprocal connectivity weights *J*_*ji*_ to have a non-zero weight *A*_*p*_. For the remaining 1−*ρ*_*pq*_ fraction of non-zero connections in the upper triangle, we set the reciprocal connectivity weights to zero. The corresponding cell-type dependent reciprocal correlations for the multi-population sparse connectivity are then

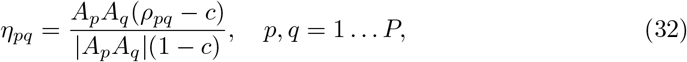

where *c* is the homogeneous connection probability (Table. 1).

**Table 1.** List of notations.

| Notation | Description |
| --- | --- |
| $i, j$ | Single neuron indices |
| $p, q$ | Neuron population indices |
| $\alpha_p$ | Fraction of neurons belonging to population $p$ |
| $\bar{J}_{pq}$ | Mean synaptic weights between populations $p, q$ |
| $J_{E/I}$ | Re-scaled mean excitatory/inhibitory synaptic weights |
| $\sigma_{z_{pq}}$ | Standard deviation of the synaptic weights between populations $p, q$ |
| $g_{pq}$ | Re-scaled standard deviation of the synaptic weights between populations $p, q$ |
| $\eta_{pq}$ | Reciprocal correlation of connectivity weights between populations $p, q$ |
| $A_{E/I}$ | Excitatory/inhibitory synaptic weights in the sparse network |
| $c$ | Homogeneous sparsity of the sparse network, $c_{pq} = c$ |
| $\sigma_{m^p}^2, \sigma_{n^p}^2$ | Variances of components on connectivity vectors $\mathbf{m}^p$ and $\mathbf{n}^p$ |
| $\sigma_{nm}^p$ | Covariance between connectivity vectors $\mathbf{m}^p$ and $\mathbf{n}^p$ |
| $\mu_x^p, \Delta_x^p$ | Population-averaged mean and variance of activation |

#### 2.1.3 Excitatory-inhibitory networks

In this work, we specifically focus on excitatory-inhibitory networks composed of *P* = 2 populations, one excitatory and one inhibitory, with respectively *N*_*E*_ and *N*_*I*_ neurons. We denote the two populations by indices *E* and *I*, so that there are four types of connections: *EE, EI, IE* and *II*. Based on the usual anatomical estimates for neocortex, we choose *N*_*E*_ = 0.8*N, N*_*I*_ = 0.2*N*, and further define *α*_*E*_ = *N*_*E*_*/N, α*_*I*_ = *N*_*I*_*/N*, as the fractions of excitatory and inhibitory neurons.

For Gaussian networks, we enforce Dale’s law only on the mean, i. e. we set 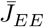 and 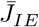 to be positive, while 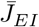 and 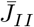 are negative. The *N*×*N* mean connectivity matrix 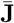 is therefore in general rank-two. To further simplify the setting, we follow [43], and consider networks where the mean weights of all excitatory connections, and respectively all inhibitory connections, are equal and set by parameters *J*_*E*_ and *J*_*I*_ :

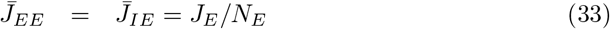

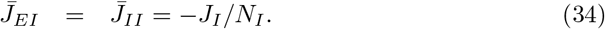

Under these additional assumptions, the entries in the first *N*_*E*_ columns of the mean connectivity matrix 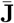 have the same positive weight *J*_*E*_*/N*_*E*_, and the entries in the following *N*_*I*_ columns have the same negative weight −*J*_*I*_*/N*_*I*_, so that 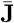 becomes rank one.

We however allow the variances 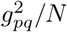 and reciprocal correlations *η*_*pq*_ to depend on both the pre- and post-synaptic population, so that the corresponding parameters form 2 × 2 matrices

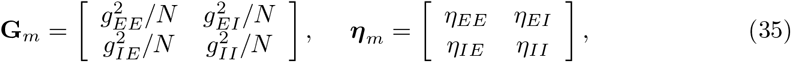

where *η*_*EI*_ = *η*_*IE*_.

For *sparse* excitatory-inhibitory networks, all non-zero excitatory (resp. inhibitory) synaptic weights are equal and positive, *A*_*E*_ *>* 0 (resp. *A*_*I*_ *<* 0). From Eqs. (27), (28) the mean and the variance of the synaptic weights in the sparse network can be matched to the parameters of the Gaussian model:

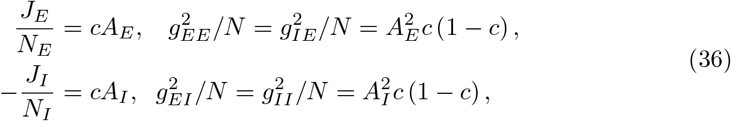

In particular, for the sparse networks with pairwise reciprocal motifs, on top of the matching means and variances, the cell-type dependent reciprocal correlations satisfy (Eq. (32))

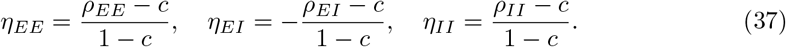

### 2.2 Globally-defined connectivity: Gaussian-mixture low-rank networks

In this section, we introduce a second broad class of connectivity models, Gaussian-mixture low-rank networks [31, 33], in which the connectivity matrix is generated from a global statistics of vectors over neurons.

Low-rank networks are a class of recurrent neural networks in which the connectivity matrix **J** is restricted to be of rank *R*, assumed to be much smaller than the number of neurons *N*. Such a connectivity matrix can be expressed as a sum of *R* unit rank terms

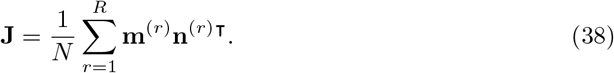

We refer to 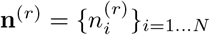 and 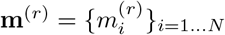 as the *r*-th left and right *connectivity vectors*. The 2*R* connectivity vectors together fully specify the connectivity matrix. Each neuron *i* is then characterized by its set of 2*R* entries 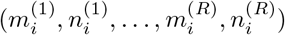 on these vectors. For unit-rank networks, the main focus of this study, the connectivity matrix is simply given by the outer product of a pair of connectivity vectors **m** and **n**:

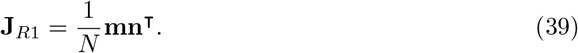

Gaussian-mixture low-rank networks are a subset of the class of low-rank networks, for which the entries of the connectivity vectors are drawn independently for each neuron from a mixture of Gaussians distribution [31]. Specifically, a fraction *α*_*p*_ of neurons is assigned to a population *p*, and within each population, the entries on the connectivity vectors are generated from a given 2*R*-dimensional Gaussian distribution. For a unit-rank network, for a neuron *i* in the population *p*, the connectivity parameters (*m*_*i*_, *n*_*i*_) are generated from a bi-variate Gaussian distribution with mean 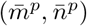, variance 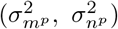 and covariance 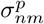.

For any unit-rank matrix of the form in Eq. (39), the only potentially non-zero eigenvalue is given by *λ* = **n**^T^**m***/N*, and the corresponding right (resp. left) eigenvector is **m** (resp. **n**). For a Gaussian-mixture model, in the large *N* limit this eigenvalue becomes

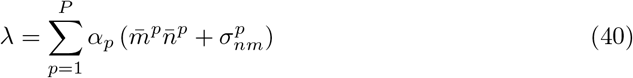

Starting from a Gaussian-mixture low-rank model in which the connectivity is globally defined, it is straightforward to compute the resulting local statistics of the connectivity, i. e. the mean 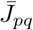, variance 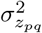 (Methods Sec. 2.1.1) and reciprocal correlation *η*_*pq*_ (Methods Sec. 2.1.2) as:

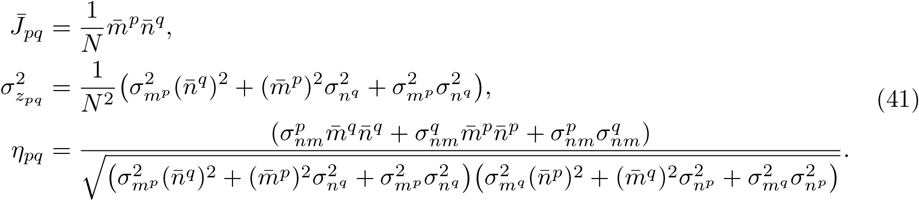

The expression for the local statistics of network connectivity using rank-*R* connectivity is in S5 Appendix.

### 2.3 Approximating locally-defined connectivity with Gaussian-mixture low-rank models

In this section, we describe our general approach for approximating an arbitrary connectivity matrix **J** with a rank-*R* matrix **J**_*R*_. We then show that for **J** corresponding to locally-defined multi-population connectivity (Methods Sec. 2.1), the resulting approximation **J**_*R*_ in general obeys Gaussian-mixture low-rank statistics as defined in Methods Sec. 2.2.

To approximate a full rank matrix **J** with a rank-*R* matrix **J**_*R*_, we use truncated eigen-decomposition, which preserves the dominant eigenvalues. We start from the full eigen-decomposition of **J**:

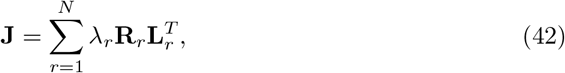

where *λ*_*r*_ is the *r*-th eigenvalue of **J** (ordered by decreasing absolute value), while **R**_*r*_ and **L**_*r*_ are the corresponding right- and left-eigenvectors that obey

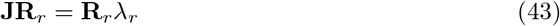

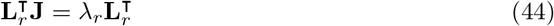

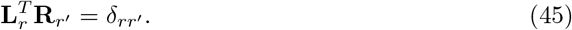

In the following, we constrain the right eigenvectors **R**_*r*_ to be of unit norm, while the normalization of the left eigenvector is determined by Eq. (45).

We obtain a rank-*R* approximation **J**_*R*_ of **J** by keeping the first *R* terms in Eq. (42):

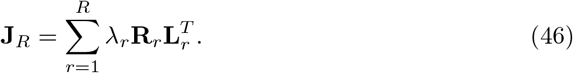

The *R* non-trivial eigenvalues and eigenvectors of **J**_*R*_ therefore correspond to the first *R* eigenvalues and eigenvectors of **J**. We then set

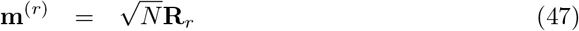

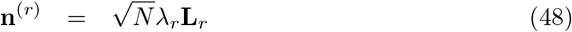

to have the same normalization for **J**_*R*_ as in Eq. (38).

To obtain a low-rank approximation for a connectivity matrix **J** generated from locally-defined statistics defined in Methods Sec. 2.1, we first determine its dominant eigenvalues and eigenvectors. Starting from Eq. (22), this problem becomes equivalent to finding the dominant eigenvalues and eigenvectors of a low-rank matrix 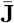 perturbed by a random matrix **Z** with block-like statistics. We compute the statistics of these eigenvalues and eigenvectors by combining the Matrix’s Determinant Lemma, the Matrix Perturbation Theory and the Central Limit Theorem. Below we summarize this general approach before applying it to different specific cases in Methods Secs. 2.4, 2.5.

We focus on the case where 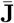 is unit rank as in the simplified excitatory-inhibitory network introduced in Methods Sec. 2.1.3. In that case, the unique non-zero eigenvalue of 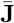 is

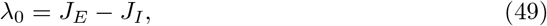

and the corresponding left and right eigenvectors are

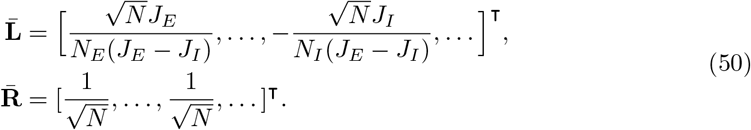

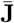 can then be rewritten as

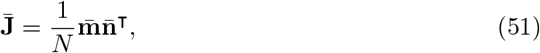

where the structure vectors 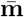 and 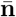 are uniquely defined by rescaling the left and right eigenvectors 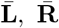 of 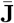 (Eq. (50)) as in Eq. (47), so that

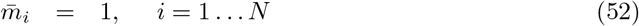

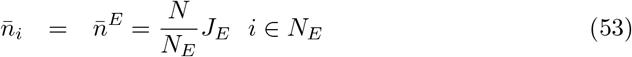

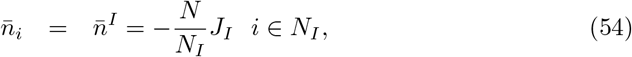

The full connectivity matrix **J** can be then expressed as

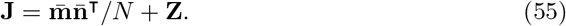

#### Eigenvalues

For a random matrix **Z** with independently distributed elements, the eigenvalues are distributed on a disk of radius *r*_*g*_ centred at the origin in the complex plane [11, 44, 50]. Correlations between elements in general modify the shape of this continuous spectrum [18, 67]. In contrast, adding a low-rank component typically induces isolated eigenvalues outside the continuous part of the spectrum [44, 45]. To obtain a low-rank approximation of the full matrix, we focus on determining these outliers when they exist.

All eigenvalues *λ* of **J** satisfy the characteristic equation

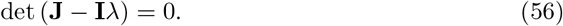

To determine the outlying eigenvalues of a random connectivity with low-rank structure, we apply the matrix determinant lemma to the l. h. s. of the characteristic equation [32]:

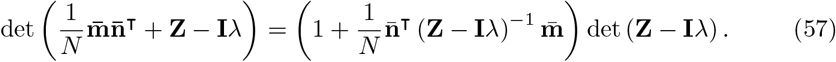

As outliers by definition cannot be eigenvalues of **Z**, they correspond to zeros of the first term in the r. h. s., and therefore satisfy:

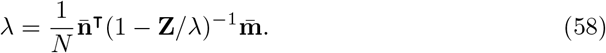

Expanding (1 − **Z***/λ*)^*−*1^ in series, we further get [32]

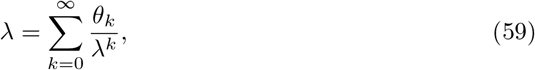

with

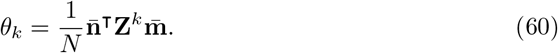

Here *θ*_0_ corresponds to the eigenvalue *λ*_0_ of 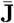 (Eq. (49)), and the higher order terms specify how this eigenvalue is modified by the random part of the connectivity. Truncating Eq. (59) at a given order, and averaging over **Z** yields a polynomial equation for the mean eigenvalues of **J**. In Methods Sec. 2.4, we exploit this equation to determine the effects of different cell-type specific random connectivity **Z** on the outlying eigenvalues.

Note that within first-order perturbation theory, the eigenvalues are given by *λ* = *λ*_0_ + Δ*λ* with

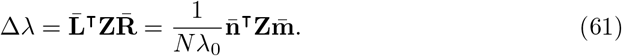

#### Eigenvectors

To determine the eigenvectors corresponding to the outlying eigenvalue of **J**, we treat it as 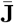 perturbed by **Z** (Eq. (22)). Matrix perturbation theory then states that, at first order, the right- and left-eigenvectors **R** and **L** of **J** corresponding to the outlying eigenvalue *λ* are given by [47]:

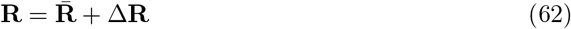

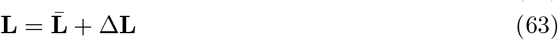

where 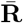 and 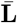 are the right- and left-eigenvectors of 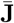 defined in Eq. (50), and

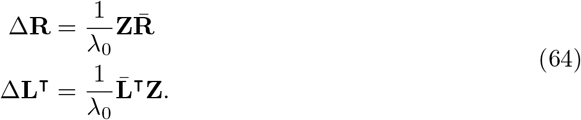

Using the normalization in Eq. (47), we then get

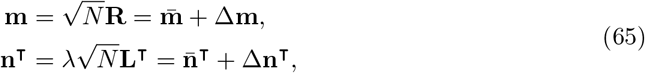

with constant entries on 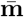 and 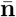 defined in Eqs. (52)-(54) and

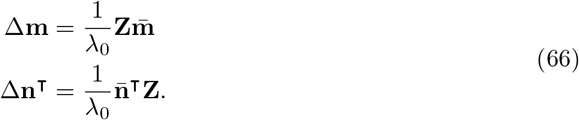

where we approximated *λ* at first order by *λ*_0_.

#### Statistics of Eigenvector entries

While 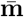 and 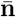 are deterministic vectors, the perturbations 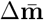 and 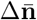 are random variables obtained by multiplying 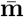 and 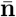 with the random matrix **Z** (Eq. (66)). We therefore next consider the statistics of the elements *m*_*i*_ and *n*_*i*_ of **m** and **n** defined in Eq. (65).

Since the elements of **Z** have zero mean, the mean values of *m*_*i*_ and *n*_*i*_ are given by 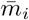 and 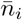 defined in Eqs. (52)-(54). The mean value of *n*_*i*_, but not *m*_*i*_, therefore depends on whether the neuron *i* belongs to the excitatory or inhibitory population. Taking into account that **Z** has block-like statistics, we split the matrix product in Eq. (66) into the sum of items corresponding to excitatory and inhibitory pre-synaptic neurons. Using Eqs. (52)-(54), Δ*m*_*i*_ and Δ*n*_*i*_ can be written as

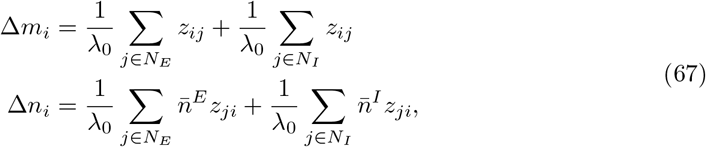

We next take the limit *N*_*E*_, *N*_*I*_→ ∞, and apply the central limit theorem, which states that each sum converges to a Gaussian random variable, so that we have

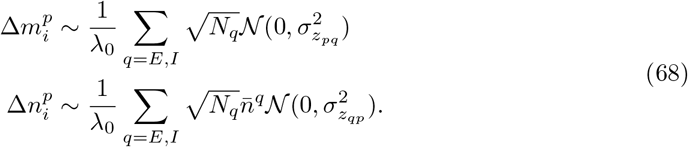

where *p* ∈ *E, I* is the population the neuron *i* belongs to, and 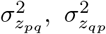 are the variance of *z*_*ij*_, *z*_*ji*_ respectively, for *i, j* in populations *p, q*. The perturbations Δ*m*_*i*_ and Δ*n*_*i*_. therefore converge to Gaussian random variables of zero mean and variances 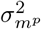 and 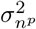 given by:

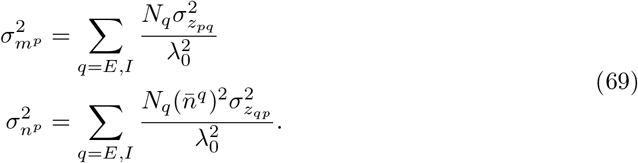

The population covariance 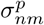 between elements *m*_*i*_ and *n*_*i*_ with *i* belonging to population *p* can furthermore be written as

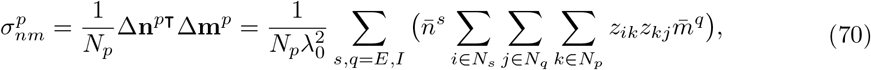

while the overall covariance *σ*_*nm*_ between all *m*_*i*_ and *n*_*i*_ reads

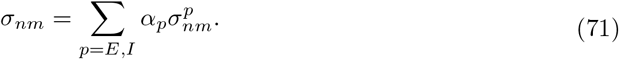

Altogether, *m*_*i*_ and *n*_*i*_ determined by perturbation theory therefore follow Gaussian-mixture statistics, where the mean and variance depend on whether the neuron *i* belongs to the excitatory or inhibitory population.

#### Comparison with simulations

The theoretical predictions for eigenvalues obtained from Eqs. (59), (60) can be verified by comparing them with the eigenvalue outliers computed by direct eigen-decomposition of the full matrix **J**. We compute the average and standard deviation of eigenvalue outliers over 30 realizations of **J**.

The predictions of perturbation theory for eigenvectors given by Eq. (64) can also be verified by direct eigen-decomposition, but to compare individual entries, an appropriate normalization is required [47]. Indeed, perturbation theory assumes that 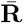 is normalized and 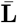 satisfies 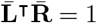 (Eq. (50)), but the perturbed eigenvectors in Eq. (62) do not obey the same normalization. We therefore first use numerical eigen-decomposition to get the right- and left-eigenvector 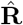 and 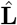 of **J**. We then normalize 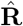 to 1, and 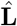 so that 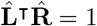. To compare 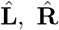 with perturbation theory, we then normalize 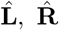 as

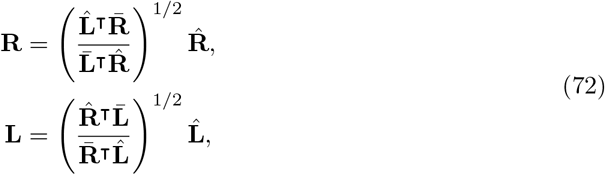

the eigenvectors **L, R** after normalization have the same statistics as 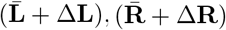 (Eqs. (50), (64)).

### 2.4 Eigenvalues

Here, we apply Eqs. (59), (60) to determine the mean and variance of outlying eigenvalues for different forms of local connectivity statistics.

#### 2.4.1 Independent random connectivity

In the case of independent random connectivity, the elements of **Z** are zero-mean, independently distributed and uncorrelated with 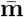 And 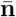. Averaging Eq. (60) over **Z** then leads to [32]:

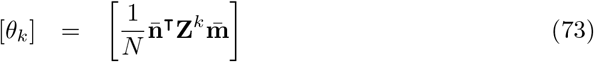

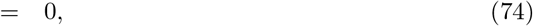

and therefore the mean eigenvalue [*λ*] of **J** is given by the eigenvalue *λ*_0_ of 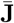. For Gaussian random connectivity, we have

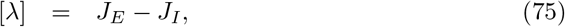

and for sparse connectivity

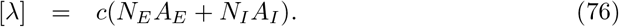

The variance 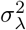 of *λ* can be computed by keeping only the linear term in Eq. (59), which leads to Eq. (61) under the assumption that *λ* ≈ *λ*_0_. Applying the central limit theorem then yields

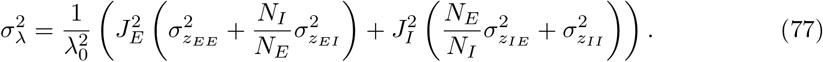

For independent Gaussian random connectivity, we substitute 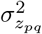 using Gaussian variance parameters (Eq. (23), (25))

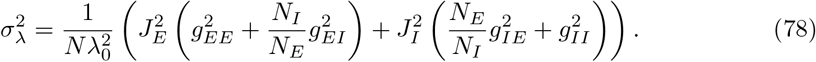

For independent sparse random connectivity, we replace 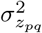 and *J*_*p*_ with the variances and means of the sparse model given in Eqs. (27), (28) and get

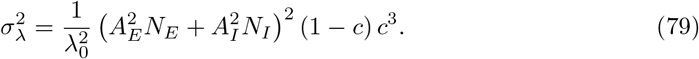

#### 2.4.2 Reciprocal motifs

In the case of connectivity with reciprocal correlations, *z*_*ij*_ and *z*_*ji*_ are correlated, so that the average [*θ*_*k*_] over **Z** in Eq. (60) is non-zero for even *k*. Here we compute [*θ*_*k*_] for *k* = 2 and truncate Eq. (59) at second order to get a third-order polynomial equation for the mean eigenvalue:

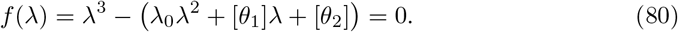

For Gaussian connectivity we have *θ*_0_ = *λ*_0_ = *J*_*E*_ − *J*_*I*_, and *θ*_1_ is given by

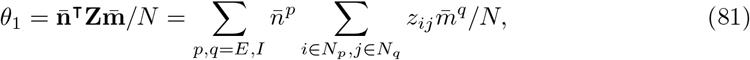

so that [*θ*_1_] = 0.

The next term *θ*_2_ is

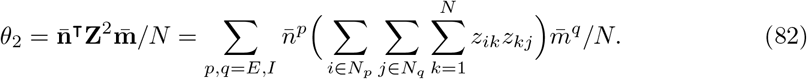

Given the reciprocal correlations defined in Eq. (29), only items with *i* = *j* in *θ*_2_ are non-zero after averaging over Gaussian realizations, so that

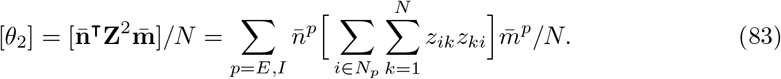

We then write

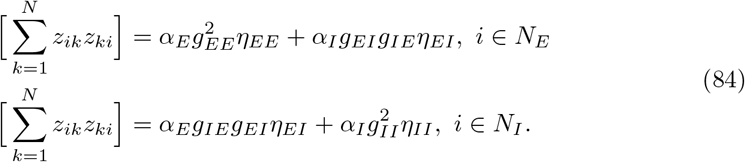

and substitute Eq. (84) and 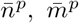 into Eq. (83) to obtain

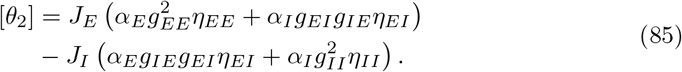

For sparse connectivity with reciprocal motifs, the correlations can be written as

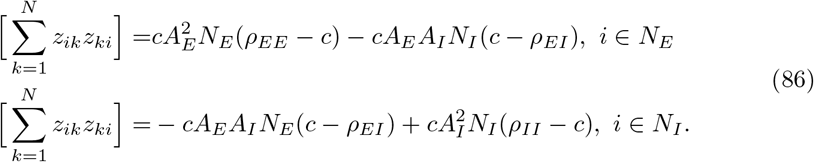

Then, combining Eqs. (36) and Eqs. (52)-(54), the second-order coefficient [*θ*_2_] for the sparse network is

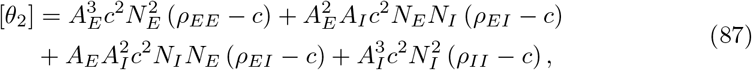

Using Eqs. (36), (37), it can be seen that Eq. (87) is equivalent to Eq. (85).

### 2.5 Eigenvectors

Here we apply Eqs. (69) and (70) to determine the variances and covariances of eigenvector entries obtained from perturbation theory for different forms of local connectivity statistics.

#### 2.5.1 Independent random connectivity

In the case of independent random connectivity, because *z*_*ik*_ and *z*_*kj*_ are not correlated in Eq. (70), the covariances 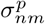 between the eigenvector entries are zero. For independent Gaussian connectivity, introducing Eq. (24) into Eqs. (69) the variances of eigenvector entries can be written as

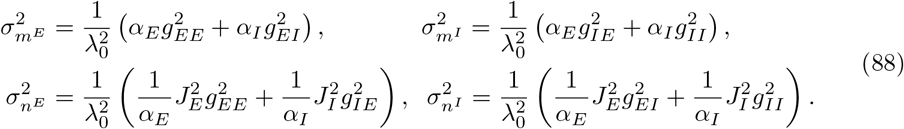

For independent sparse connectivity, substituting Eq. (28) into Eqs. (69), leads to

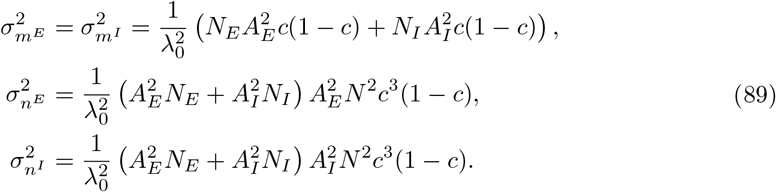

#### 2.5.2 Reciprocal motifs

In the case of connectivity with reciprocal correlations, the variances of eigenvector entries are identical to the independent case.

As we have shown in Eq. (70), noise correlation between the rank-one vectors arises from the correlation between pairwise random connectivity weights in the situation with reciprocal motifs, only items with *i* = *j* (for *z*_*ik*_, *z*_*kj*_) in the same population *q* are non-zero, so that we have

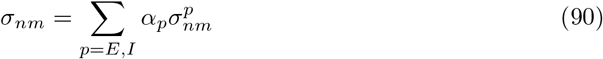

with

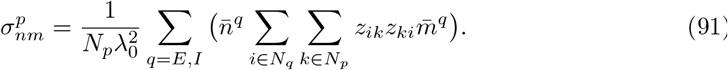

For Gaussian connectivity with reciprocal correlations, the covariances between entries on **Z** matched by population can be written as

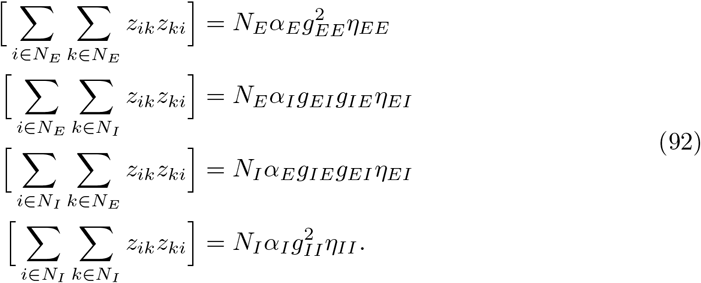

Combining Eq. (92) and the mean rank-one connectivity loadings Eqs. (52)-(54), we obtain the population covariances as

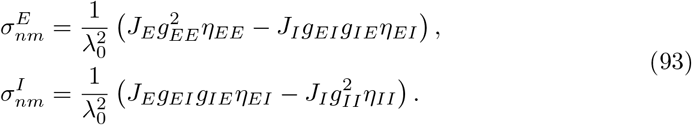

We note that the large deviation of the dominant eigenvalue *λ* in the network with reciprocal motifs also increases the nonlinearity of the vector perturbations. To account for this nonlinearity, we start from Eq. (40) for *λ* and get 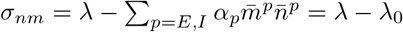, then we compare with Eq. (80) and get the approximation relationship

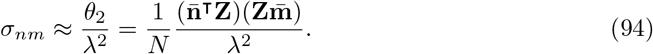

Similarly, for the covariance of each population we have

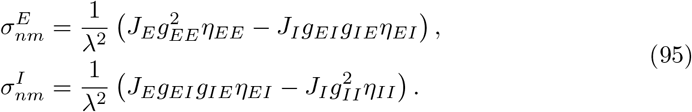

For sparse connectivity with reciprocal correlations, the calculations are similar, with entries of **Z** being Bernoulli-distributed

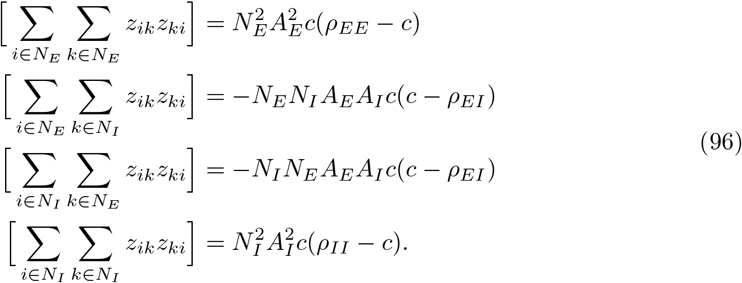

and we have the population covariance

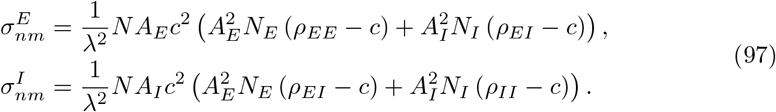

Using Eqs. (36), (37), it can be seen that Eq. (97) is equivalent to Eq. (95).

### 2.6 Dynamics

In this section, we show how approximating locally-defined connectivity by a global low-rank structure allows us to analyse the emerging low-dimensional dynamics. We first summarize the mean-field theory (MFT) for Gaussian-mixture low-rank networks [31, 33]. We then apply it to unit-rank connectivity obtained as an approximation of locally-defined connectivity. We finally compare the resulting description of the dynamics with an alternate mean-field approach for random connectivity consisting of a superposition of low-rank and full-rank random parts as in Eq. (22) [30, 32].

Throughout this study, we consider recurrent networks of rate units with recurrent interactions defined by a connectivity matrix **J**. The dynamical activity of unit *i* is represented by a variable *x*_*i*_(*t*), which we interpret as the total synaptic input current. The firing rate of unit *i* is given by *r*_*i*_(*t*) = *ϕ*(*x*_*i*_(*t*)) where *ϕ*(*x*) = 1 + tanh (*x* − *θ*) is a positive transfer function. We focus on networks without external inputs, so that the dynamics of synaptic input to neuron *i* is given by

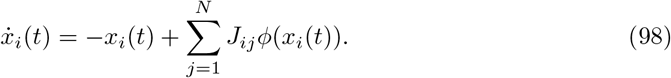

In Figs 5-7, we compare the dynamics determined by direct simulations for a locally-defined connectivity matrix with a mean-field description obtained for a unit-rank approximation.

#### 2.6.1 Mean-field theory for Gaussian-mixture low-rank connectivity

Here we review the mean-field theory for networks in which the connectivity matrix is exactly low-rank, with components of connectivity vectors moreover drawn from Gaussian-mixture distribution. Previous works have shown that in this case, the dynamics of the collective activity **x**(*t*) = {*x*_*i*_}_*i*=1…*N*_ are embedded in a linear subspace of dimension *R* spanned by the connectivity vectors **m**^(*r*)^ [30–33]. Thus, **x**(*t*) can be expressed as

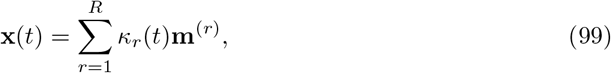

where *κ*_*r*_(*t*) for *r* = 1 … *R* are collective latent variables that quantify the components of **x**(*t*) along the connectivity vectors **m**^(*r*)^. We assume that **m**^(*r*)^ are orthogonal to each other, so that *κ*_*r*_(*t*) can be expressed as

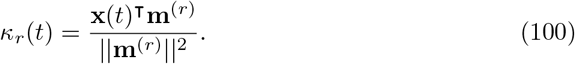

For simplicity, here we moreover assume that the initial value of **x**(*t*) lies in the subspace spanned by the vectors **m**^(*r*)^. More generally, the initial state can be included as an additional input to the dynamics [31, 33].

For a unit rank connectivity **J** = **mn**^⊤^*/N*, there is a single latent variable *κ* corresponding to the connectivity vector **m**, and the dynamics of **x**(*t*) is expressed as

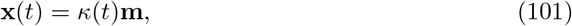

with *κ*(*t*) given by

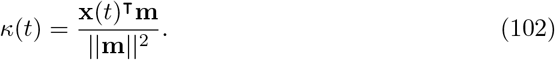

Substituting Eq. (101) into Eq. (98) and inserting the unit-rank connectivity, the dynamics of the latent variable *κ* can be expressed as

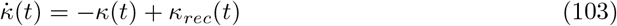

where

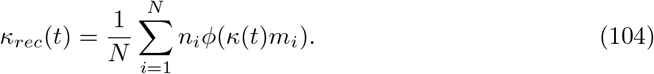

The quantity *κ*_*rec*_(*t*) represents the total recurrent input to *κ*. The sum in the r. h. s. of Eq. (104) can moreover be interpreted as the empirical average of *n*_*i*_*ϕ*(*κ*(*t*)*m*_*i*_) over the neurons in the network. In the limit of large network size *N*, this average converges to the integral of *nϕ*(*κ*(*t*)*m*) over the distribution *P* (*m, n*) of the components of connectivity vectors. For low-rank networks, the mean-field limit corresponds to replacing *κ*_*rec*_(*t*) with this integral [31, 33]:

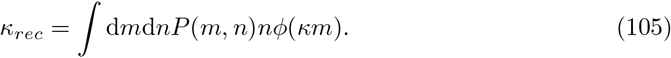

In the Gaussian-mixture low-rank model, each neuron *i* is assigned to a population *p* for *p* = 1 … *P*. Within each population, the components (*m*_*i*_, *n*_*i*_) are generated from a multivariate Gaussian distribution *P*^*p*^(*m, n*), that is

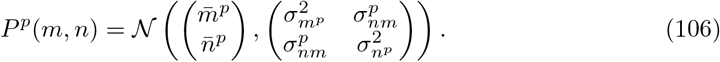

In the mean-field limit, *κ*_*rec*_ is therefore given by

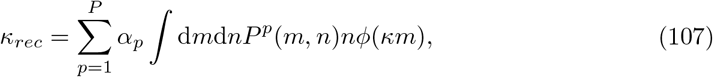

where α_*p*_ is the fraction of neurons in population *p*.

Integrating by parts, *κ*_*rec*_ can be re-expressed as (S1 Appendix)

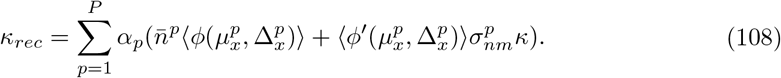

Here 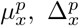 are the mean and variance of the inputs to population *p*, given by

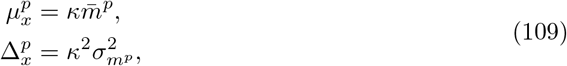

and the symbol ⟨*f* (*μ*, Δ)⟩ stands for the expected value of a function *f* (*x*) with respect to a Gaussian variable *x* with mean and variance *μ*, Δ, that is

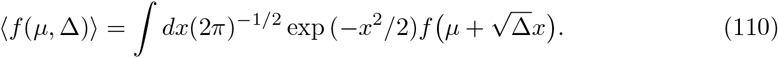

Altogether, using MFT for Gaussian-mixture low-rank networks gives the closed dynamics of the latent variable *κ*:

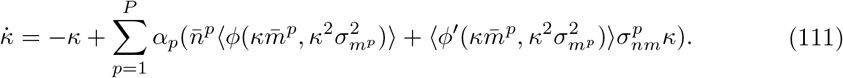

In particular, the corresponding steady state is given by

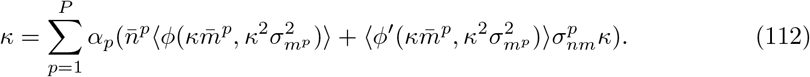

Note that the first and second terms on the r. h. s. respectively correspond to the mean and covariance of the entries of the unit-rank connectivity vectors **m** and **n**.

#### 2.6.2 Approximate dynamics for locally-defined connectivity

We next apply the MFT to unit-rank connectivity obtained as an approximation of locally-defined connectivity for the different considered cases.

##### Independent connectivity

We start from the network with independent connectivity, in which case the unit-rank connectivity vectors obtained by approximating locally-defined connectivity have no covariance, i. e. 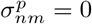 (Methods Sec. 2.5).

The dynamical system for the latent variable *κ* therefore contains only the mean term

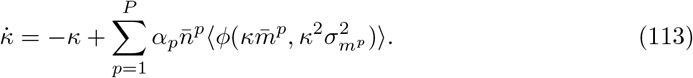

For the Gaussian random model, inserting the expressions for 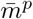 and 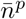 (Eqs. (52)-(54)), the fixed point obeys

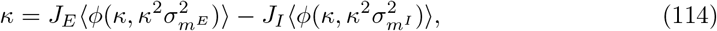

where the variance 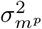 of connectivity components *m*_*i*_ is given by Eq. (88).

For the sparse random model, we further consider 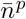 given by Eqs. (36), (52)-(54) and the fixed point is

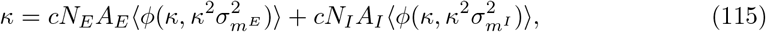

where 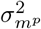 is obtained from Eq. (89).

##### Reciprocal motifs

Correlations between reciprocal connections lead to non-zero covariance 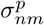 between the unit-rank connectivity vectors obtained by approximating locally-defined connectivity (Methods Sec. 2.3, Eq. (70)). The dynamical system for the latent variable *κ* therefore contains both the mean and covariance terms (Eq. (112)).

For the Gaussian random model, combining Eqs. (52)-(54), (88), (95) the fixed point obeys

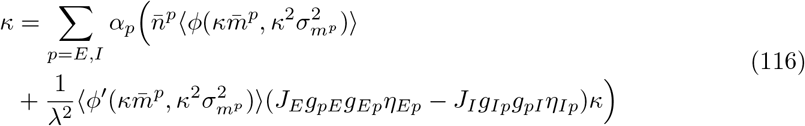

with the variance 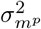 of connectivity components *m*_*i*_ given by Eq. (88). For the sparse model, combining Eqs. (36), (52)-(54), (89), (97), the fixed point obeys

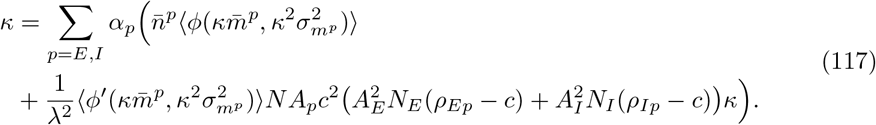

with the variance 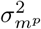 of connectivity components *m*_*i*_ given by Eq. (89).

#### 2.6.3 Mean-field theory for superpositions of low-rank and full rank random connectivity

Here we review an alternate form of mean-field theory for random connectivity consisting of a superposition of a low-rank structure and full-rank random part [30, 32]. This form of MFT can be directly applied to independently generated connections, where the connectivity matrix consists precisely of a superposition of a low-rank part corresponding to the mean, and a full-rank random part corresponding to fluctuations (Eqs. (22), (119)). Extending this type of MFT to the situation where reciprocal connections are present is however challenging [18]. Moreover, in contrast to the case where connectivity is exactly low-rank, when the additional full-rank random part is present the mean-field theory describes only the steady-state activity (and linearized dynamics around it), but not the full dynamics as in Eq. (98).

The key assumption of MFT for randomly connected networks is that the total input *x*_*i*_ to each unit can be approximated as a stochastic Gaussian process [52]. The first two cumulants (mean and variance) of that Gaussian process are then computed self-consistently to characterize the steady-state activity.

At a fixed point, the total input *x*_*i*_ obeys

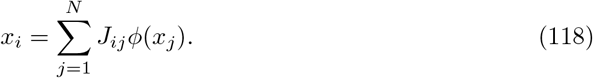

Replacing *J*_*ij*_, where *i, j* belong to populations *p, q* respectively, by the superposition of rank-one mean and full-rank random connectivity components 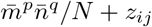 we get

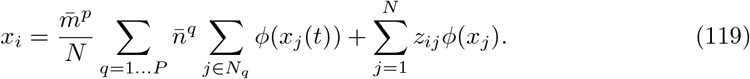

Denoting by [·] the average over the distribution of *x*_*i*_, the mean of *x*_*i*_ can then be expressed as

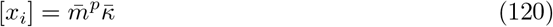

where we introduced

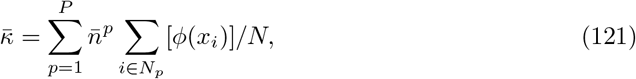

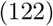

and we assumed that the zero-mean random connectivity *z*_*ij*_ is uncorrelated with the firing rate *ϕ*(*x*_*j*_), so that

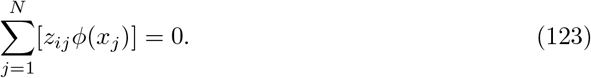

Similarly, the correlation between *x*_*i*_ and *x*_*j*_, where *i* ∈ *N*_*p*_ and *j* ∈ *N*_*q*_, is given by

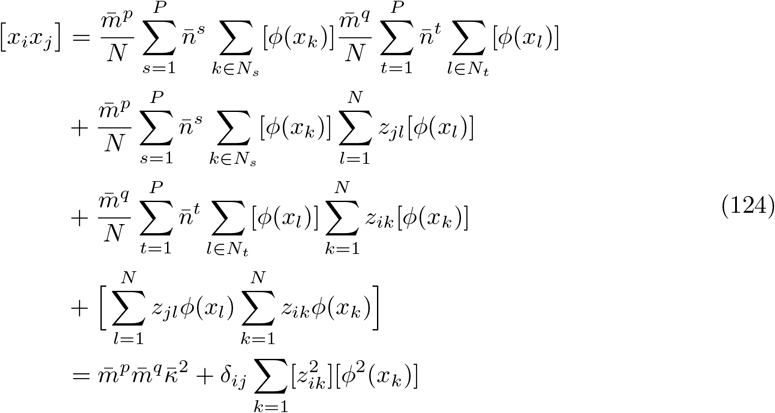

where we assume the neuronal activities are decorrelated [*ϕ*(*x*_*i*_)*ϕ*(*x*_*j*_)] = [*ϕ*(*x*_*i*_)][*ϕ*(*x*_*j*_)] when *i* ≠ *j*. This assumption holds for independently-generated connections, but not in presence of reciprocal correlations [18]. The covariance between *x*_*i*_ and *x*_*j*_ therefore becomes

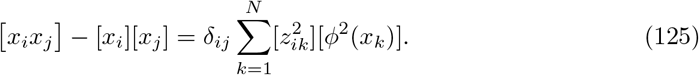

Within the mean-field approximation, neuronal activation *x*_*i*_ are therefore uncorrelated Gaussian variables with mean and variance given by Eqs. 120 and 125

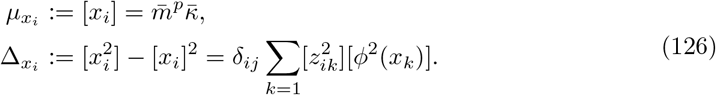

To determine 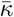 and [*ϕ*(*x*_*k*_)^2^], we finally express Eqs. (121) and (126) as Gaussian integrals over *x*_*i*_ in population *p*:

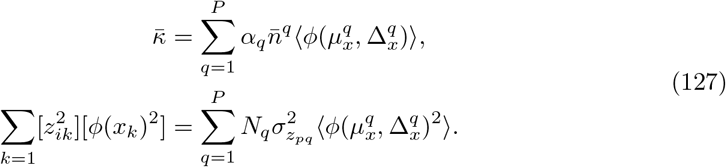

Here we replaced 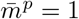 and 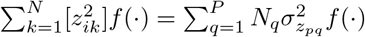 given the eigenvector normalization in Eq. (52), and the assumption that variances 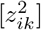 depend on the populations the units *i* and *k* belong to (Eqs. (24), (28), (52)). Therefore, the stationary mean and variance of the dynamics of synaptic inputs in population *p* are

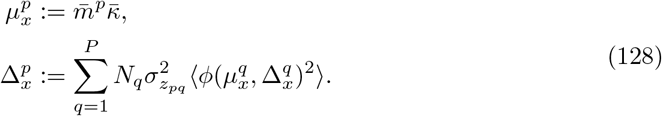

Eqs. (128), (127) give the self-consistent equations for the stationary solutions of the dynamics.

More specifically, in the Gaussian random model, we combine connectivity statistics given by Eqs. (24), (52)-(54), so that we have

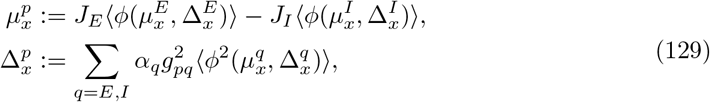

while for the sparse random model, we combine connectivity statistics given by Eqs. (27), (28), (36), (52)-(54), so that we have

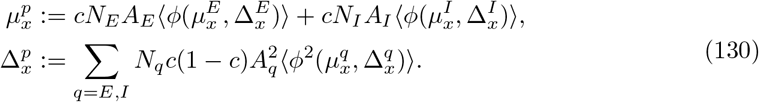

## Code Availability

Code will be made available upon publication.

## Supporting information

### S1 Appendix. Dynamics in Gaussian-mixture low-rank networks

Here, we provide the derivation for the dynamics of the latent variable *κ* (Eq. (111)) in the Gaussian-mixture low-rank network model. We consider a rank-one connectivity consisting of *P* populations, with the neurons in each population accounting for a *α*_*p*_ percentage of all neurons. The entries on the left and right eigenvectors **n**^*p*^, **m**^*p*^ assigned to population *p* are sampled from a multivariate Gaussian distribution with mean 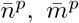, variance 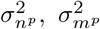 and covariance 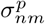 (Methods Sec. 2.3) So, *κ*_*rec*_ in Eq. (107) is further decomposed into two integrals involving the contributions from the mean and random connected components, respectively,

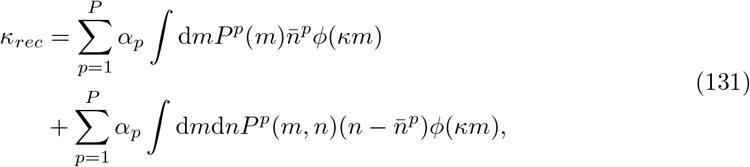

here, *P*^*p*^(*m*) represents the marginal Gaussian distribution for connectivity loadings *m*_*i*_ of neurons in population *p*

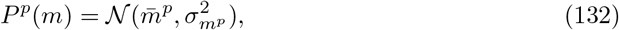

and *P*^*p*^(*m, n*) represents the multivariate Gaussian distribution for connectivity loadings *m*_*i*_, *n*_*i*_ of neurons in population *p* (Eq. (106)).

Note that the synaptic input to neurons in population *p* is a scaled Gaussian variable *x*_*i*_ = *κm*_*i*_ corresponding to the Gaussian loadings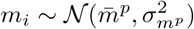, so it conforms to a Gaussian distribution 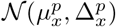 with the population-averaged mean and variance

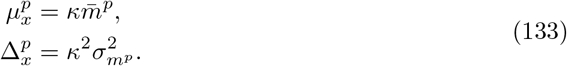

Thus, the first term in Eq. (131) is

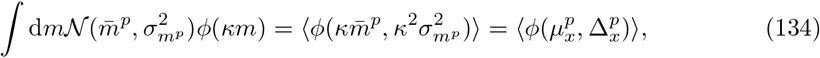

a Gaussian integral term.

We then use Stein’s lemma

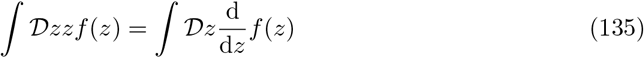

and further replace the multivariate Gaussian distribution by Eq. (106), to compute the second term in Eq. (131) attributed to the random connectivity component as

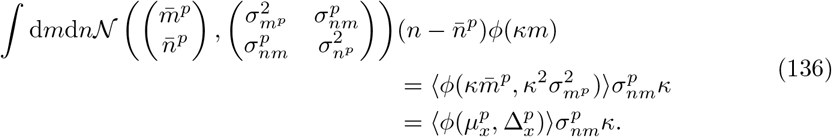

Finally, combining the contributions from both the mean and random components, we re-express *κ*_*rec*_ in Eq. (107) and retrieve Eq. (111).

### S2 Appendix. Linear stability at fixed points in rank-one networks

To determine the stability of fixed points of the network with rank-one connectivity structure, we consider the rank-one connectivity **J**_*R*1_ and study the stability of the one-dimensional latent dynamical variable *κ* in the neighbourhood area of its fixed point *κ*^0^. We set the fixed point of synaptic input 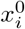, define the perturbation of the latent variable *κ*^1^ and the perturbation of the synaptic input 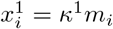, the temporal evolution of *κ*^1^ is expressed as

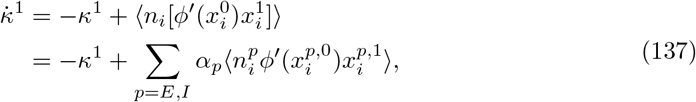

for the rank-one approximation network **J**_*R*1_ = **mn**^⊤^*/N*, we remove symbol [·]. Next, in the Gaussian-mixture low-rank framework, the entries 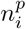 and 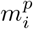 are jointly sampled from a bivariate Gaussian distribution characterized by means, variances and covariances, 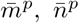 and 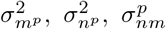. Using a similar approach to previous studies [30, 31], we set

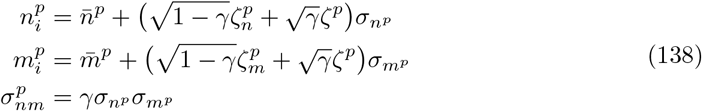

where 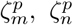 and 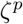 are three independent normal random variables 𝒩(0, 1). By substituting variables in Eq. (138), after some linear algebra, we get

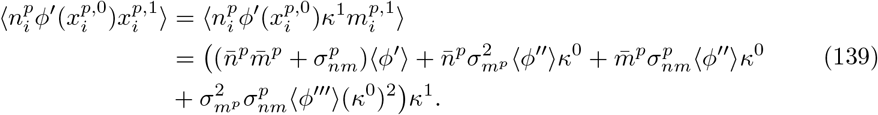

We finally obtain the time evolution of *κ*

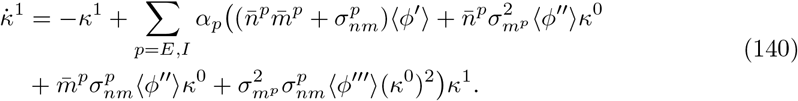

The Jacobian for the latent dynamical variable’s fixed point *κ*^0^ is

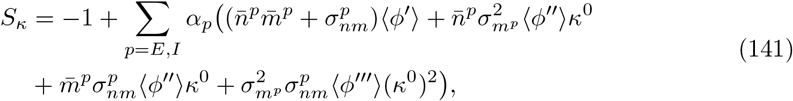

which is a scalar in our rank-one network. The fixed point is stable when *S*_*κ*_ *<* 0 and unstable when *S*_*κ*_ *>* 0. All the above parameters are expressed and calculated in the text, so that we can check the stability of each fixation point using the formulas above.

### S3 Appendix. Comparison between dynamics in full-rank connectivity with rank-one approximation

Here, we compare the dynamics solved in the full-rank network with dynamics solved by the rank-one approximation, considering the limitations of applying the classical MFT, we only discuss networks with independent connections.

Using the rank-one approximation, global dynamics is characterized by the latent dynamical variable *κ*, which satisfies

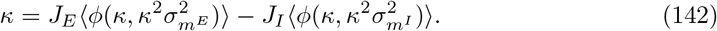

Combining the connectivity statistics given in Eqs. (52)-(54), (88), we thus express the self-consistent equations for the population mean and variance of the synaptic inputs *x*_*i*_ using *κ* (Eq. (109))

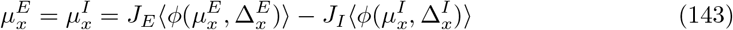

and

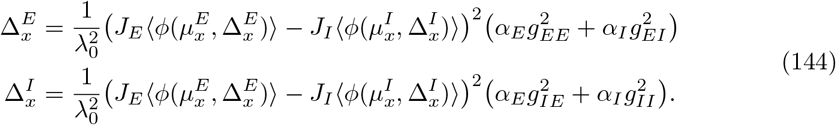

Comparing with the self-consistent equations for population mean and variance in the full-rank network Eq. (129), we find in particular that the expression for 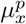 is the same in both representations, but the expressions for the variance 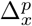 are different. Specifically, the low-rank approximation shows that the different heterogeneity between excitatory and inhibitory populations depends only on the block-structured variances, i. e., 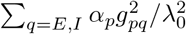, and independent of the historical population activity. The full-rank dynamics, on the other hand, shows that the different heterogeneity depends on the local relationship between block structured variance and the structure of historical population activity [68].

For a simplified network example, where the locally defined connectivity has homogeneous random parameters *g*_*pq*_ = *g, p* = *E, I*, the variance of the rank-one perturbation eigenvector 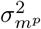 is thereby the same for both excitatory and inhibitory populations. Because *λ*_0_ = *J*_*E*_ − *J*_*I*_, means and variances in Eqs. (143), (144) are

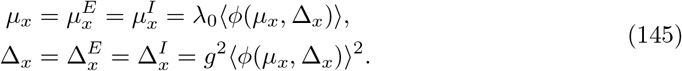

So that, for this simple example, the heterogeneity of dynamics in the full-rank approximation is *g*^2^⟨(*ϕ*(*μ*_*x*_, Δ_*x*_))^2^⟩ (Eq. (129)), while the heterogeneity in the rank-one approximation is *g*^2^⟨*ϕ*(*μ*_*x*_, Δ_*x*_)⟩^2^, the difference does not substantially change the bistable transition and the performance of the low-rank approximation.

### S4 Appendix. Chaotic dynamical transition point for networks with independent connectivity

Conventionally, the collective dynamics of random neural network becomes chaotic when the effect of random component is too strong. Based on previous study [11], when the effective random gain, also known as the radius of the random eigenvalue bulk *r*_*g*_, exceed the critical threshold 1, the network enters the chaotic dynamical state.

Considering the block-like E-I random network with i. i. d. random connectivity, we define a ℝ^2*×*2^ matrix with elements 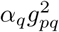

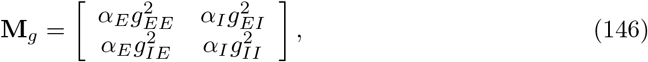

the first eigenvalue of **M**_*g*_ determines the radius of the continuous eigenvalues bulk of **J**, that is

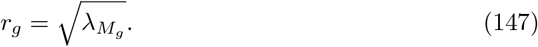

For the sparse E-I network, considering the relationships given by Eq. (36), we calculate the radius for the sparse network as

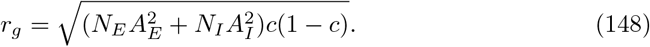

### S5 Appendix. Local connectivity statistics in rank-*R* Gaussian-mixture models

We show that the statistical properties of the entries on the rank-*R* connectivity vectors 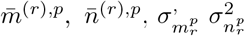 and 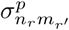 directly determine the means 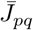, variances 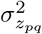 and reciprocal correlations *η*_*pq*_ of the resulting local synaptic weights *J*_*ij*_, where *i* ∈ *N*_*p*_, *j* ∈ *N*_*q*_. Considering Eq. (2), the cell-type-dependent mean

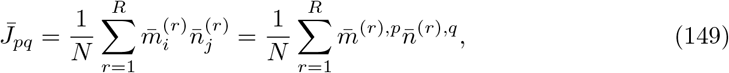

and the cell-type-dependent variance of locally defined connections is

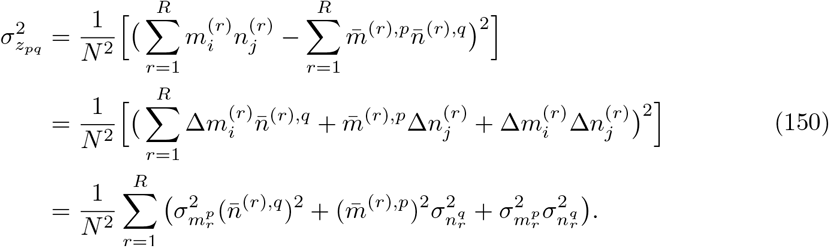

Finally the correlation between the pairwise weights *J*_*ij*_, *J*_*ji*_ is computed as

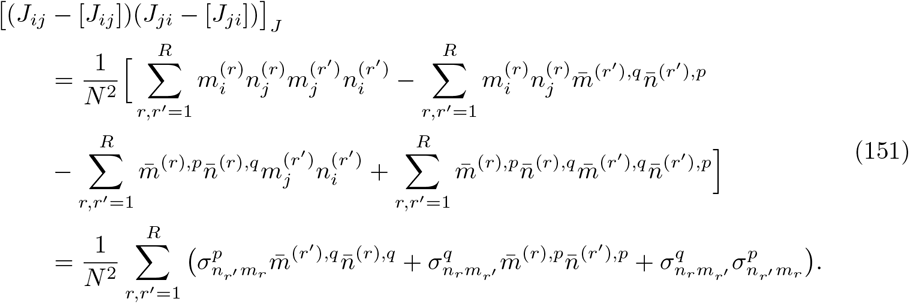

Substituting Eqs. (150), (151) into Eq. (29) leads to the resulting reciprocal correlation *η*_*pq*_.

## Acknowledgments

The project was supported by the Eranet-Neuron project IMBALANCE and the program “Ecoles Universitaires de Recherche” launched by the French Government and implemented by the ANR, with the reference ANR-17-EURE-0017. There are no competing interests.

